# SALLB not targeted by IMiDs is important for maintenance of liver cancer cells

**DOI:** 10.1101/2023.07.07.548071

**Authors:** Kim Anh L. Vu, Shiva Moein, Kalpana Kumari, Bee Hui Liu, Chong Gao, Miao Liu, Meng-Yuan Dai, Zhaoji Liu, Feng Li, Jing Ping Tang, Danilo Maddalo, Douglas S. Auld, Dominick E. Casalena, Xi Tian, Mahmoud A. Bassal, Viktoriia Iakovleva, Migara Kavishka Jayasinghe, Justin L. Tan, Alicia J. Stein, Christopher Budjan, Claire Xinyu Shao, Qiling Zhou, Patrick D. Fischer, Logan H. Sigua, Jun Qi, Haribabu Arthanari, Gerburg M. Wulf, Daniel G. Tenen, Li Chai

## Abstract

Immunomodulatory (IMiD) drugs have shown a prominent therapeutic activity in hematologic malignancies; however, their usage in solid tumors is limited. The oncofetal protein SALL4 is essential for cancer cell survival. While IMiDs can induce SALL4 degradation, they fail to induce cell death in SALL4-expressing cancer cell lines. Here, we observed that this inefficacy arose from their selective degradation of the long SALL4 isoform, while sparing the short SALL4B isoform. Selective silencing of SALL4B phenocopied total SALL4 depletion by inducing cancer apoptosis, underscoring the critical role of SALL4B in cancer maintenance. Recognizing that IMiDs can’t degrade SALL4B, we performed a high-throughput screen to identify compound(s) that could achieve this. We identified a small molecule compound that degrades both SALL4 isoforms with enhanced potency towards SALL4B in a cereblon- and proteasome-dependent manner. This compound suppressed cancer cell proliferation and attenuated tumor development in both cell line and patient-derived xenograft models. Transcriptomic analyses further revealed convergent effects of genetic and pharmacologic SALL4B depletion on DNA damage response and replication pathways. Together, these findings identify SALL4B as the therapeutically relevant isoform in SALL4-dependent cancers and establish isoform-aware targeted degradation as a strategy to overcome the limitation of IMiDs in solid tumors.

**Graphical Abstract | Identification of QE:** a non-IMiDs degrader capable of degrading both SALL4A and SALL4B, triggers anti-cancer effects beyond IMiDs, and Impacts of QE on Key Validated SALL4B Targets and Pathways

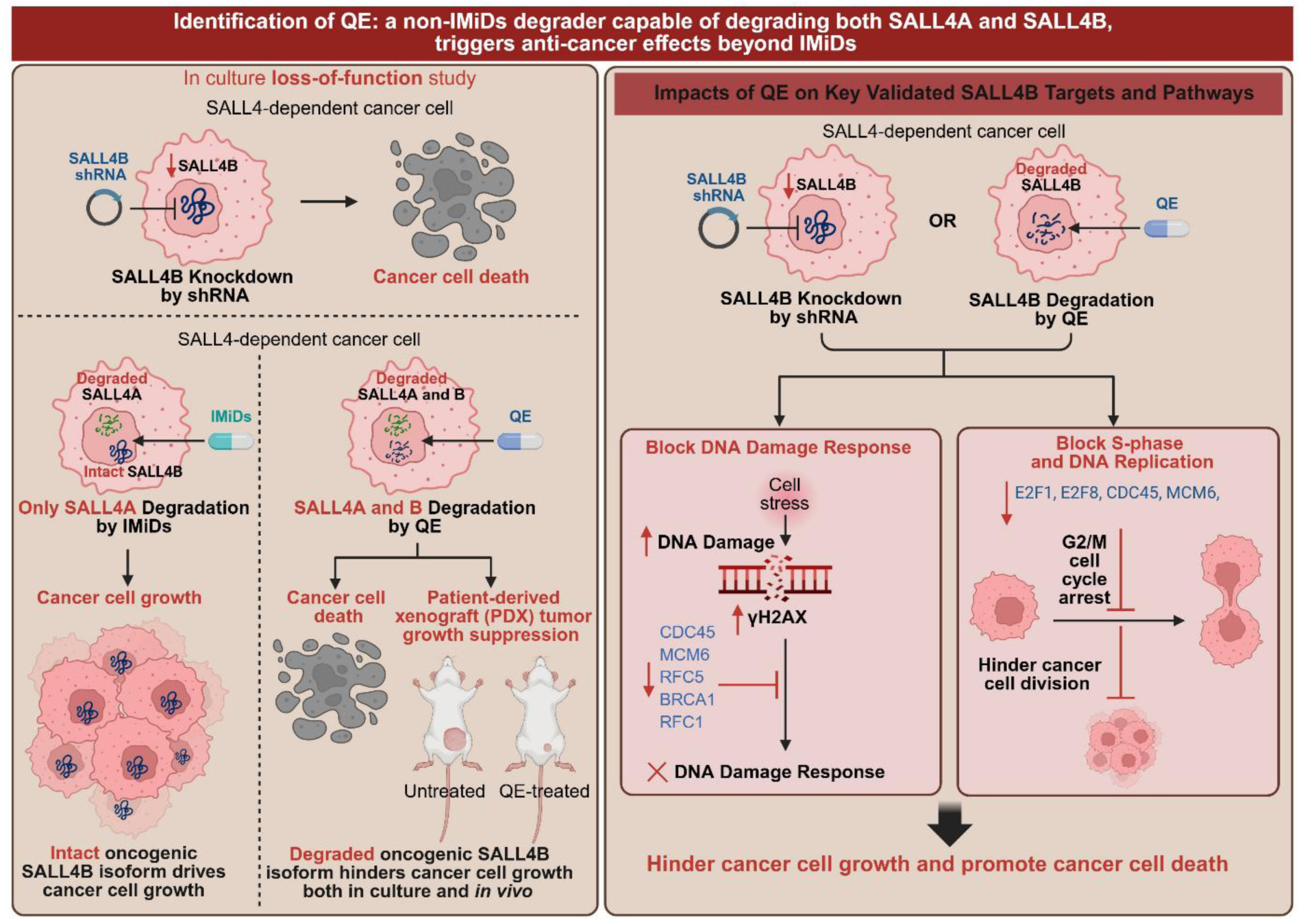

## Introduction

Targeted protein degradation is an emerging drug discovery technology that aims to chemically knockdown pathological proteins by hijacking the cell’s proteolysis machinery(1). This strategy is particularly useful for targeting currently undruggable protein targets(2), such as transcription factors. The oncofetal protein SALL4 (Spalt-like transcription factor 4) is a C2H2 zinc finger transcription factor expressed during embryonic development, where it regulates genes responsible for stemness, self-renewal, migration, and anti-apoptosis(3, 4). In most adult human tissue, SALL4 is silenced, but it is re-expressed and up-regulated in many human cancers, especially upon hypomethylating regimens (5, 6). High SALL4 level correlates with more aggressive disease, metastasis, and poor overall survival in HCC(4, 7), lung(8), and endometrial(9) patients, indicating its value as a prognostic marker(10, 11). Our group and others have shown that downregulating SALL4 expression using short hairpin RNA interference in human and xenotransplant murine models of HCC, endometrial cancer, myeloid leukemia, and gastric cancer leads to potent anti-proliferative effect and tumor growth attenuation.(4, 7, 9, 12, 13) These findings demonstrate the critical role of SALL4 in driving tumorigenesis and cancer cell survival, making it a promising therapeutic target for cancer treatment. Since SALL4 is largely silenced in adult tissues and reactivated only in cancer, pharmacologic degradation of SALL4 is expected to have minimal tissue toxicity in adult cancer treatment.

Currently, immunomodulatory drugs (IMiDs) are reported to induce CRBN-mediated SALL4 degradation(14, 15). However, in this study, we found that IMiDs only degrade the SALL4A isoform and fail to inhibit the growth of SALL4-positive cancer cells. We discovered that SALL4B, the isoform not affected by IMiDs, is required for the survival of SALL4-mediated cancer cells. These new insights guided the discovery of a small molecule degrader that induces proteasome-dependent degradation of both SALL4A and SALL4B isoforms. This compound potently suppressed tumor growth in cell line xenografts as well as patient-derived xenograft (PDX) HCC models, without major side effects, demonstrating strong therapeutic potential.

RNA-seq-based gene expression profiling revealed a shared transcriptional signature between genetic knockdown and pharmacologic degradation of SALL4B in HCC cells, characterized by alterations in DNA damage response, S-phase, and DNA replication pathways. These findings uncover mechanistic dependencies in SALL4-driven cancers and support dual SALL4 isoform degradation as a therapeutic strategy.

## Results

### Degradation of SALL4A by IMiDs does not affect cancer cell survival

Given that SALL4 is essential for cancer cell survival, and IMiDs have been reported to degrade SALL4(14, 15), we investigated whether IMiDs (Fig. 1a) could be used to treat SALL4-driven cancers. SALL4 has two major isoforms, SALL4A and SALL4B, generated through alternative splicing. The longer isoform, SALL4A, contains four zinc finger clusters (ZFC1, ZFC2, ZFC3 and ZFC4), whereas the shorter isoform, SALL4B, only has ZFC1 and ZFC4 (Fig. 1b). Surprisingly, we found that SALL4-mediated cancer cell lines SNU-398 and H661 were insensitive to IMiDs treatment (Fig. 1c, d), similar to SALL4-negative controls SNU-387 and SNU-423 (Fig. 1e). Notably, IMiDs only degraded SALL4A, but not SALL4B, in SNU-398 cells (Fig. 1f and Supplementary Fig. 1a, b), without affecting SALL4 mRNA levels (Fig. 1g). This specific SALL4A degradation by IMiDs was confirmed using an isoform-specific luciferase reporter system^28^ in H1299 cells overexpressing SALL4A or SALLB (Fig. 1h and Supplementary Fig. c,d). We further found that deletion of ZFC2 rendered SALL4A resistant to pomalidomide-induced degradation, whereas deletion of ZFC1, ZFC3, or ZNF4 did not alter its sensitivity (Fig. 1i,j). Consistent with the absence of ZFC2, SALL4B was not degraded by pomalidomide (Fig. 1j).

**Figure 1.**
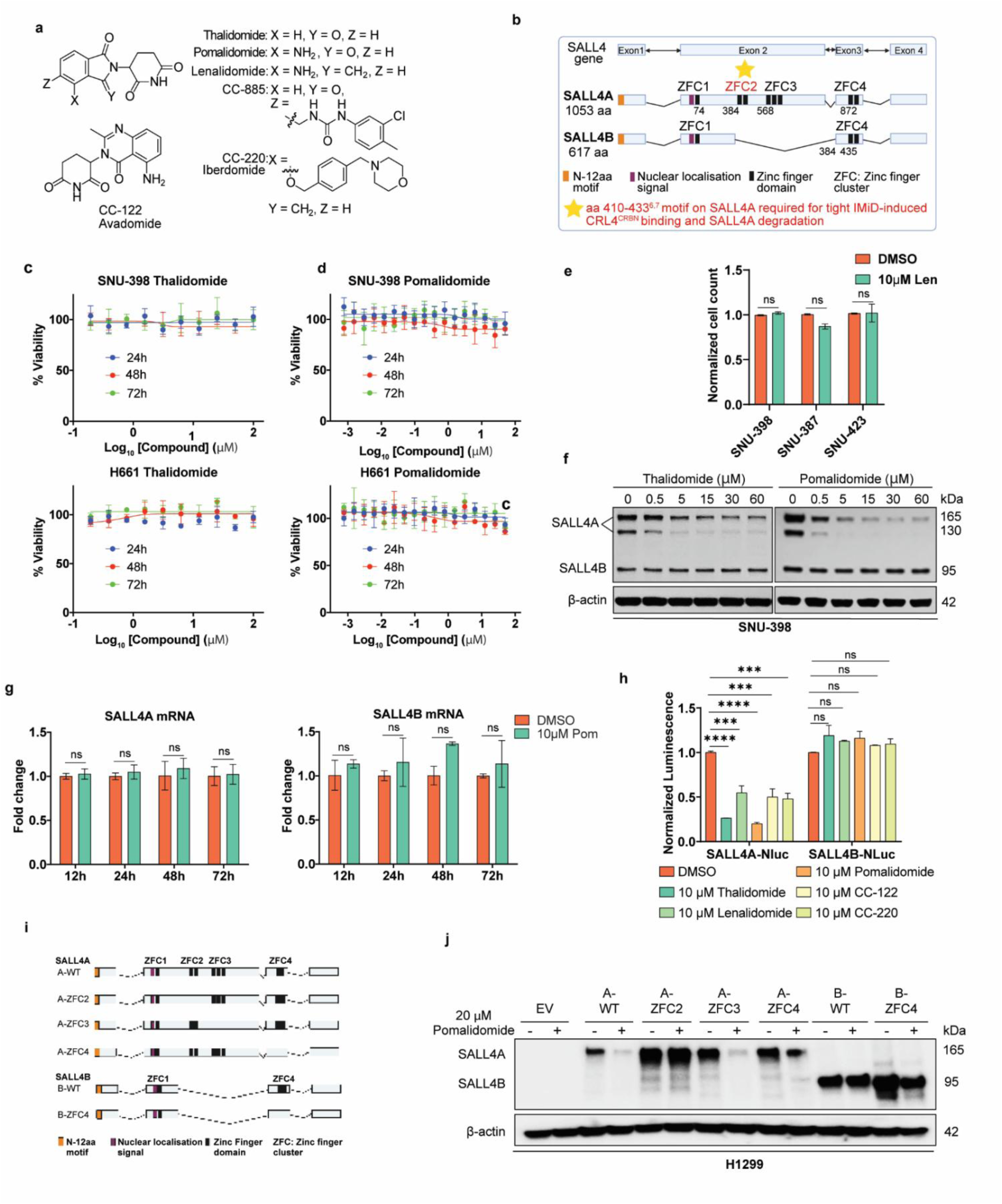
The SALL4B isoform drives oncogenesis. **a, Chemical structures of IMiDs.** b, Schematic of SALL4 isoforms. c,d, Viability of SNU-398 and H661 cells treated with IMiDs (thalidomide, pomalidomide) for 24, 48 and 72 hours (a, n = 3; b, n = 4; mean ± SD). e, Viability of SNU-398, SNU-387 and SNU-423 cells after 72 hours of treatment with lenalidomide (10 µM) (n = 2; mean ± SD). f, Immunoblot showing IMiDs selectively degrade SALL4A but not SALL4B in SNU-398 cells (8 hours, representative of n = 2). g, qPCR analysis of SALL4 mRNA in SNU-398 cells treated with pomalidomide for the indicated times (n = 3; mean ± SD). h. Nluc/Fluc ratios after 16 hours IMiD treatment in H1299 cells expressing SALL4A-Nluc or SALL4B-Nluc (n = 2; mean ± SD). i, Schematic of H1299 isogenic cell lines expressing wild-type (WT) or zinc-finger-deleted SALL4A or SALL4B. j, Immunoblots showing resistance of SALL4B and ZFC2-deleted SALL4A to pomalidomide-induced degradation after 24 hours treatment in H1299 isogenic lines described in (i). Statistics: Unpaired two-tailed t-test for (c) and (f). One-way ANOVA with Dunnett’s post hoc test versus DMSO for (h). *P < 0.05, **P < 0.01, ***P < 0.001, ****P < 0.0001; ns, not significant (P ≥ 0.05).

Collectively, these findings show that degradation of SALL4A alone does not impair cancer cell survival, implying a dependency on SALL4B or compensation by SALL4B in SALL4-driven cancers.

### Downregulation of SALL4B is essential for inhibiting SALL4-mediated cancer cell survival

To investigate the importance of SALL4B for cancer survival, we designed an shRNA targeting the unique region of SALL4B mRNA to selectively silence SALL4B (Fig. 2a). We then compared the effect of SALL4B-specific knockdown on cancer cell apoptosis and survival to that of total SALL4 knockdown. In SALL4-high SNU-398 HCC cells, SALL4B knockdown increased apoptosis by ∼21.9% (measured by Annexin V staining), comparable to total SALL4 knockdown (Fig. 2b, Supplementary Fig. 2a). SALL4B silencing alone similarly reduced SNU-398 cell viability (Fig. 2c) and impaired survival and growth of SNU-398 in both soft agar colony formation (Fig. 2d, Supplementary Fig. 2b) and clonogenic assays (Fig. 2g). Consistent results were observed in additional SALL4-high HCC and lung cancer cells (Fig. 2e,f,h). In contrast, SALL4-low cancer cells were unaffected by either total SALL4 or SALL4B-specific knockdown (Fig. 2i). Together, these data demonstrate SALL4B as the critical isoform supporting the survival of SALL4-dependent cancer cells.

**Figure 2.**
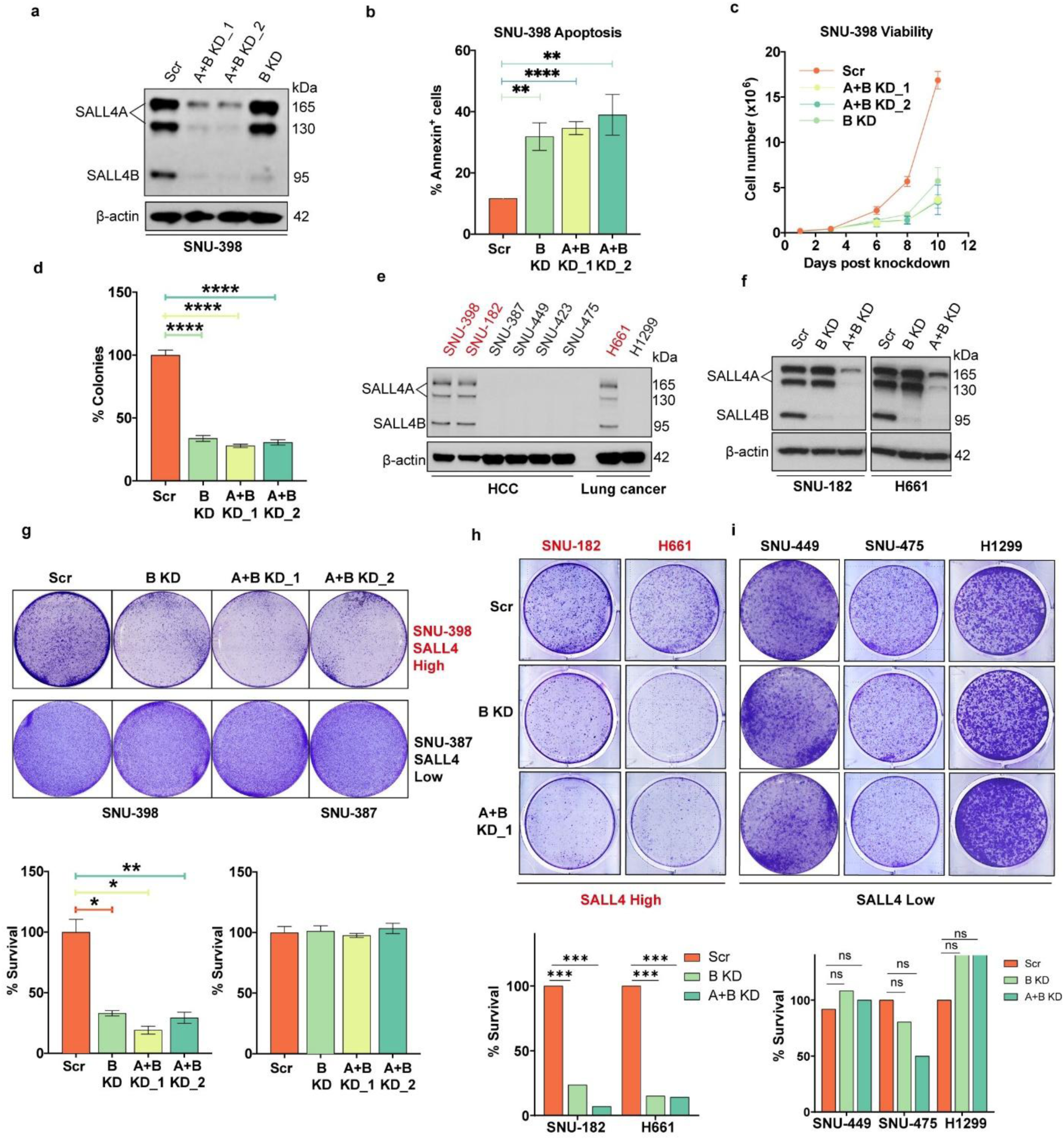
**a, Immunoblot showing total SALL4 and SALL4B-specific knockdown in SNU-398 cells.** b, Annexin V+ SNU-398 cells 5 days after Scr or SALL4 shRNA transduction (n = 3; mean ± SD). c, Growth curves of SNU-398 cells post-transduction with SALL4 shRNA or Scr (n = 3; mean ± SD). d, Soft agar colony formation of SNU-398 cells following Scr or SALL4 shRNA (n = 5; mean ± SD). e, Immunoblots showing differential SALL4 expression across HCC and lung cancer cell lines. f, Total SALL4 and SALL4B-specific knockdown in SNU-182 (HCC) and H661 (lung) cells. g, Clonogenic assay images (top) and quantification (bottom) for SALL4-high SNU-398 and SALL4-low SNU-387 cells transduced with Scr or SALL4 shRNA (n = 3; mean ± SD). h,i, Representative clonogenic assay images (top) and quantification (bottom) for SALL4-high (h) and SALL4-low (i) HCC and lung cancer cells infected with Scr or SALL4 shRNA. Statistics: Two-way repeated-measures ANOVA with Dunnett’s post hoc test vs Scr for (f); one-way ANOVA with Dunnett’s post hoc test vs Scr for (e,g,h,k,l). *P < 0.05; **P < 0.01; ***P < 0.001; ****P < 0.0001; ns, not significant.

To assess the role of SALL4B in the initiation and maintenance of HCC *in vivo*, we generated and studied SALL4B transgenic mice(16). While none of the 19 wild-type controls developed liver tumors (Supplementary Fig. 2c, panel i, ii), 50% (7 of 14) of aged SALL4B transgenic mice (>16 months) developed spontaneous HCC (Supplementary Fig. 2c, iii, iv; Supplementary Table 1). To assess whether young SALL4B transgenic mice exhibit increased susceptibility to liver tumor development, we employed a two-stage chemical carcinogenesis protocol using N-nitrosodiethylamine (DEN) and Phenobarbital (PB) to induce pre-neoplastic lesions. Mice were examined 10 and 23 weeks after PB treatment (Supplementary Fig. 2d). Consistently, young SALL4B transgenic mice displayed increased mitotic activity and a higher incidence of hepatic lesions following carcinogen exposure (Supplementary Fig. 2e, iv-viii, Supplementary Table 2&3). These findings support a role for SALL4B in promoting hepatocarcinogenesis *in vivo*.

### Identification of QE, a non-IMiDs chemical probe capable of degrading SALL4B

We then initiated a high-throughput screening of 50,000 small molecules to identify compounds that could bind and degrade both SALL4 isoforms, specially SALL4B, for the treatment of SALL4-mediated cancer. The screening cascade included thermal shift assays, SALL4B luciferase reporter assays, immunoblotting and cell viability assays (Fig. 3a). The thermal shift assay identified 883 potential binders to the 1-300aa N-terminal domain of SALL4, which is shared by both SALL4A and SALL4B. From these, 18 hits were identified as potential SALL4B degraders via the SALL4B luciferase reporter assay. Immunoblotting confirmed that 5 of these hits (Supplementary Fig. 3a) effectively degraded endogenous SALL4A and SALL4B (34.7 to 81.8%) in SNU-398 HCC cells after 16h of treatment at 15 μM (Supplementary Fig. 3b). Among these 5 hits, QE was chosen as the lead compound due to its greater potency in SALL4-high compared to SALL4-low cancer cells, demonstrating up to 58-fold differential sensitivity, with 72h EC_50_ values of 350 nM in SNU-398 cells and 880 nM in H661 cells (Supplementary Fig. 3c, d).

**Figure 3.**
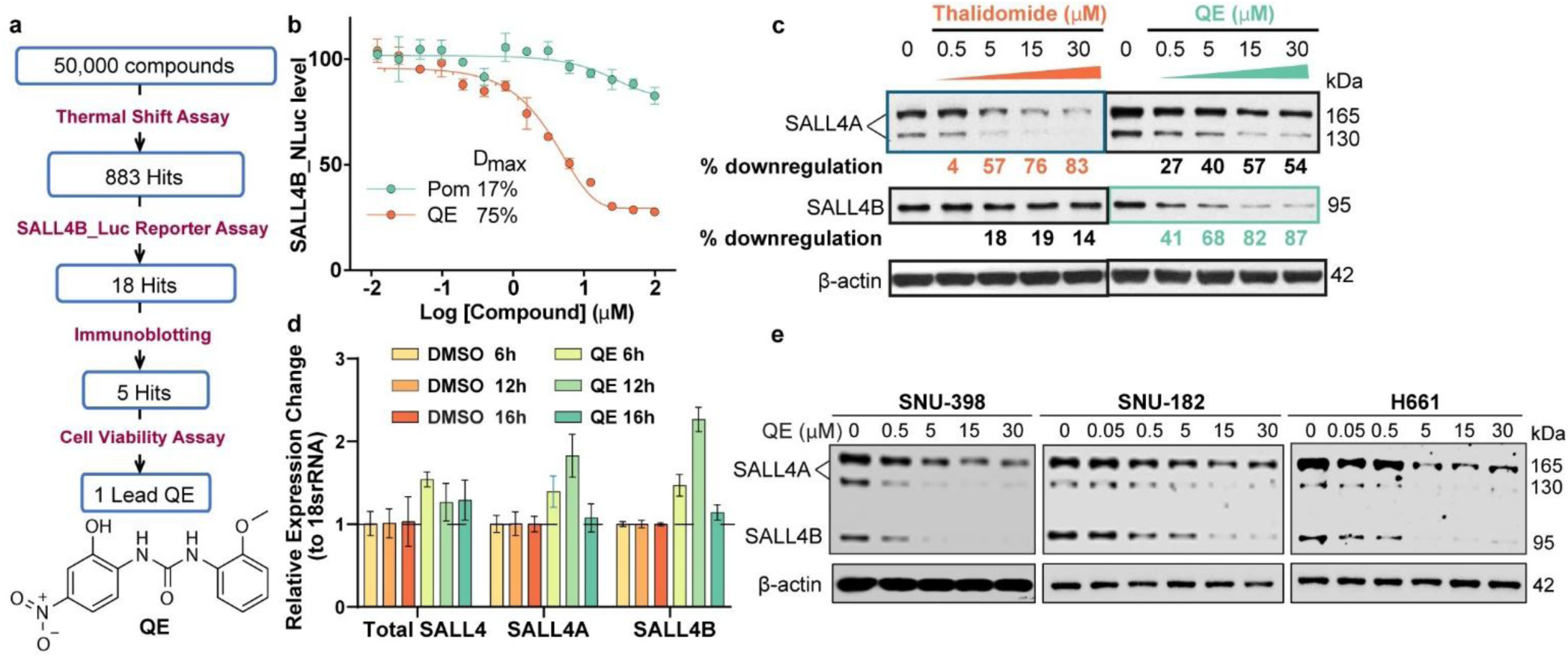
QE, but not IMiDs, induces SALL4B degradation. **a,** High-throughput screening identifying QE inducing SALL4B degradation. **b,** QE, but not Pomalidomide, induced dose-proportionate degradation of SALL4B-luciferase in H1299 cells after 9 hours (n = 3, mean ± SD). **c**, Immunoblot showing QE, but not Thalidomide, induced dose-dependent degradation of endogenous SALL4B in SNU-398 cells after 18 hours. **d,** qRT-PCR results showing total SALL4, SALL4A and SALL4B mRNA levels in SNU-398 cells were not affected by QE treatment (5 µM) at all time points (n = 3, mean ± SD). **e,** Immunoblots showing endogenous SALL4B degradation after 18h QE treatment in SNU-398, SNU-182 and H661 cells (representative of n = 2). **Statistics:** Two-way ANOVA with Šídák’s multiple comparisons test for (d) revealed no significant effect of treatment or treatment × time interaction; no post-hoc comparisons reached significance (ns).

To compare the activity of QE with that of IMiDs, we evaluated isoform-specific degradation and associated cellular consequences using an isoform-specific SALL4 luciferase reporter system (Supplementary Fig. 1c) and SNU-398 HCC cells that express high endogenous SALL4. H1299 cells overexpressing SALL4B_Nluc and SNU-398 cells were treated with increasing concentrations of QE or IMiDs. Following treatment, SALL4B_Nluc levels were measured via luciferase assay, and endogenous SALL4A and B proteins were assessed by immunoblotting.

QE induced a pronounced dose-dependent reduction of SALL4B protein in both the reporter assay (maximal degradation D_max_ = 75%; Fig. 3b) and SNU-398 cells (D_max_ = 87%; Fig. 3c). In contrast, IMiDs (pomalidomide or thalidomide) exhibited a limited effect on the SALL4B_Nluc (D_max_ = 17%; Fig. 3b) or endogenous SALL4B (D_max_ = 19%; Fig. 3c), indicating that SALL4B degradation is achieved only with QE treatment.

In SNU-398 cells, QE potently degraded SALL4B (DC_50_ ∼ 0.64 μM, D_max_ = 87%) but was less effective at degrading SALL4A (D_max_ = 57%, DC_50_ ∼ 4.23 μM). Conversely, IMiDs preferentially degraded SALL4A (D_max_ = 83%, DC_50_ = 7.38 μM) but showed minimal activity toward SALL4B (D_max_ = 19%, DC_50_ > 30 μM) (Fig. 2c). Importantly, QE degraded both SALL4A and SALL4B without affecting SALL4 mRNA levels, indicating a post-translational mechanism (Fig. 3d).

QE also induced dose-proportional degradation of endogenous SALL4A and SALL4B across multiple SALL4-mediated cancer cell lines, including SNU-398 and SNU-182 HCC cells, and H661 lung cancer cells (Fig. 3e).

Collectively, these findings demonstrate that QE, but not IMiDs, effectively degrades both SALL4 isoforms, including the oncogenic SALL4B.

### The therapeutic effect of QE depends on the depletion of SALL4, especially SALL4B

To get an insight about the stoichiometry of QE and SALL4 interaction, the binding of QE to purified SALL4(1-300aa) N-terminal domain was assessed. SALL4(1-300aa) was incubated with increasing concentrations of QE, and protein stabilization was measured by changes in protein melting temperature (ΔTm). QE induced a dose-dependent increase in SALL4(1-300aa) ΔTm, indicating that QE bound and stabilized SALL4(1-300aa). Notably, when the molar ratio of QE to SALL4(1-300aa) approached approximately 1:1, ΔTm reached a plateau (Fig. 4a). This suggests that QE engages the 1 to 300 N-terminal domain of the SALL4 protein, which is present in both SALL4A and SALL4B isoforms, via a stoichiometric 1:1 binding.

**Figure 4.**
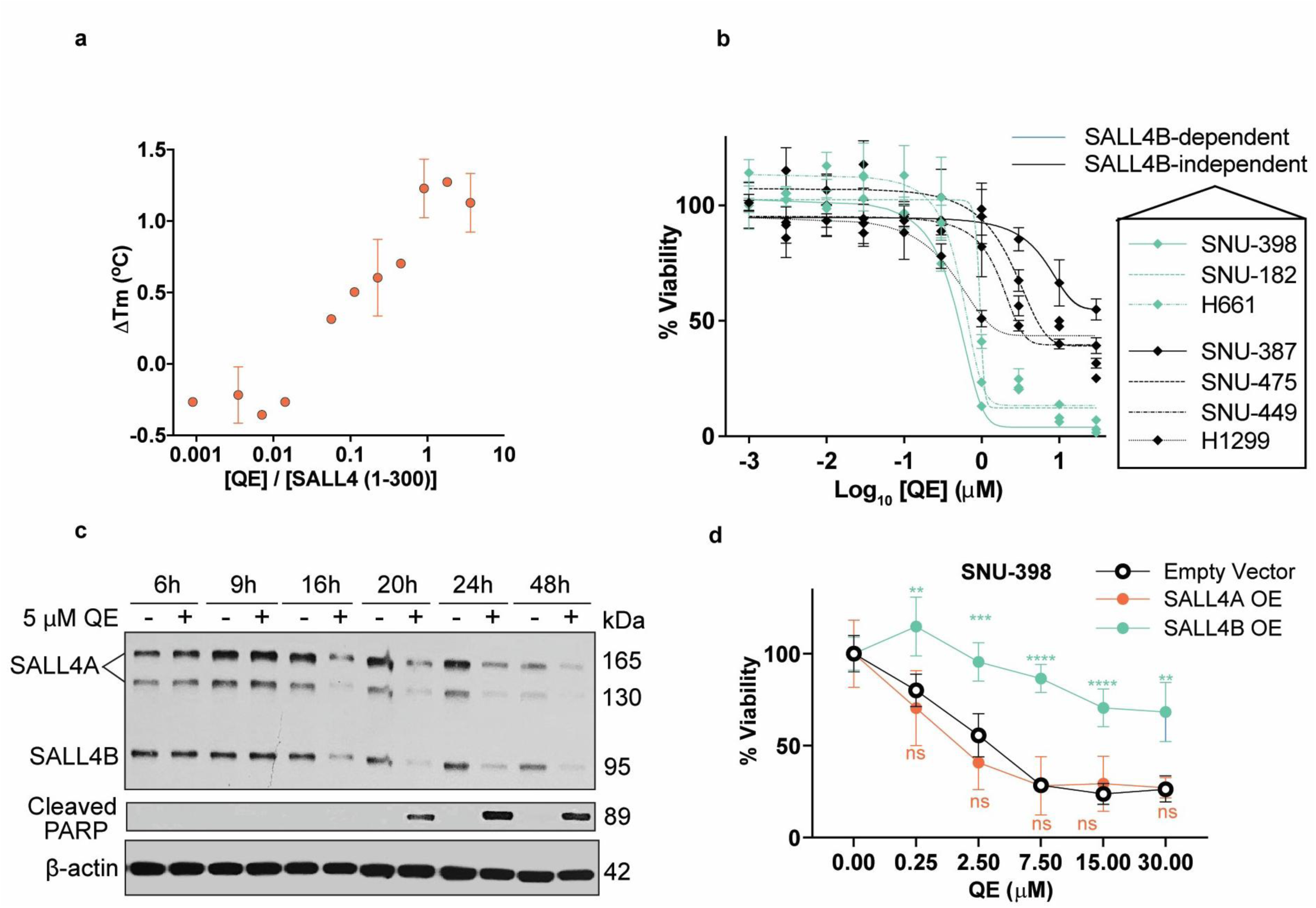
QE causes enhanced loss of viability in SALL4B-dependent HCC cells. **a,** SALL4(1-300aa) thermal stabilization by QE after 30-min incubation (n = 2, mean ± SD). **b,** Viability of SALL4B-dependent (SNU-398, SNU-182, H661) and SALL4B-independent (SNU-387, SNU-475, SNU-449. H1299) HCC and lung cancer cells after 72h QE treatment (n = 3, mean ± SD). **c,** Immunoblot showing kinetics of SALL4B degradation and PARP cleavage induced by 5 µM QE in SNU-398 cells (representative of n = 2). **d,** Overexpression of SALL4B, but not SALL4A, conferred resistance to QE (24h treatment) in SNU-398 cells (n = 6, mean ± SD). Two-way ANOVA with Dunnett’s post-hoc test for (d) comparing SALL4A OE and SALL4B OE to Empty Vector. **P < 0.01, ***P < 0.001, ****P < 0.0001; ns, not significant (P > 0.05).

We next assessed the cellular response to QE across a panel of HCC and non-small cell lung carcinoma (NSCLC) cell lines with differential dependency on SALL4 for survival. Following 72 h of QE treatment, SALL4-dependent lines, including SNU-398, SNU-182, and H661 (Fig. 2e), exhibited greater sensitivity, achieving near-complete growth inhibition (E_max_ = 90-100%) at 1μM with nanomolar IC_50_ values of 397nM, 951nM, and 522nM, respectively (Fig. 4b). After 20 h of QE exposure, these cells also showed strong apoptotic responses marked by increased levels of the apoptotic marker cleaved PARP (Fig. 4c). In contrast, SALL4-independent lines (SNU-387, SNU-449, SNU-475, and H1299; Fig. 1h, l) were less sensitive, with 30-60% of cells remaining viable even at 30 μM QE (Fig. 4b). Mechanistically, QE treatment resulted in degradation of SALL4 in sensitive cells (Fig. 2e, h).

To further interrogate isoform-specific contributions, we overexpressed SALL4A or SALL4B in SNU-398 cells. SALL4B overexpression conferred marked resistance to QE, whereas SALL4A overexpression or empty vector controls remained sensitive to QE treatment (Fig. 4d, Supplementary Fig. 4a, b).

Together, these results indicate that the cytotoxic activity of QE in SALL4-driven cancers is linked to its ability to deplete SALL4, with SALL4B playing a role in determining QE sensitivity.

### QE promotes proteasomal SALL4B degradation via cereblon

We then investigated the pathway through which QE induces SALL4 degradation. In SNU-398, SNU-182, and H661 cells, QE-mediated SALL4 degradation was markedly reduced by the proteasome inhibitor MG132 and by MLN4924, an inhibitor of cullin-RING E3 ligase activation (Fig. 5a). Correspondingly, QE-induced apoptosis was substantially diminished in the presence of proteasome inhibitors (Fig. 5b), and SALL4-dependent cells displayed increased viability when co-treated with these inhibitors, whereas SALL4-independent SNU-387 cells showed no protective effect (Fig. 5c, Supplementary Fig. 5a).

**Figure 5.**
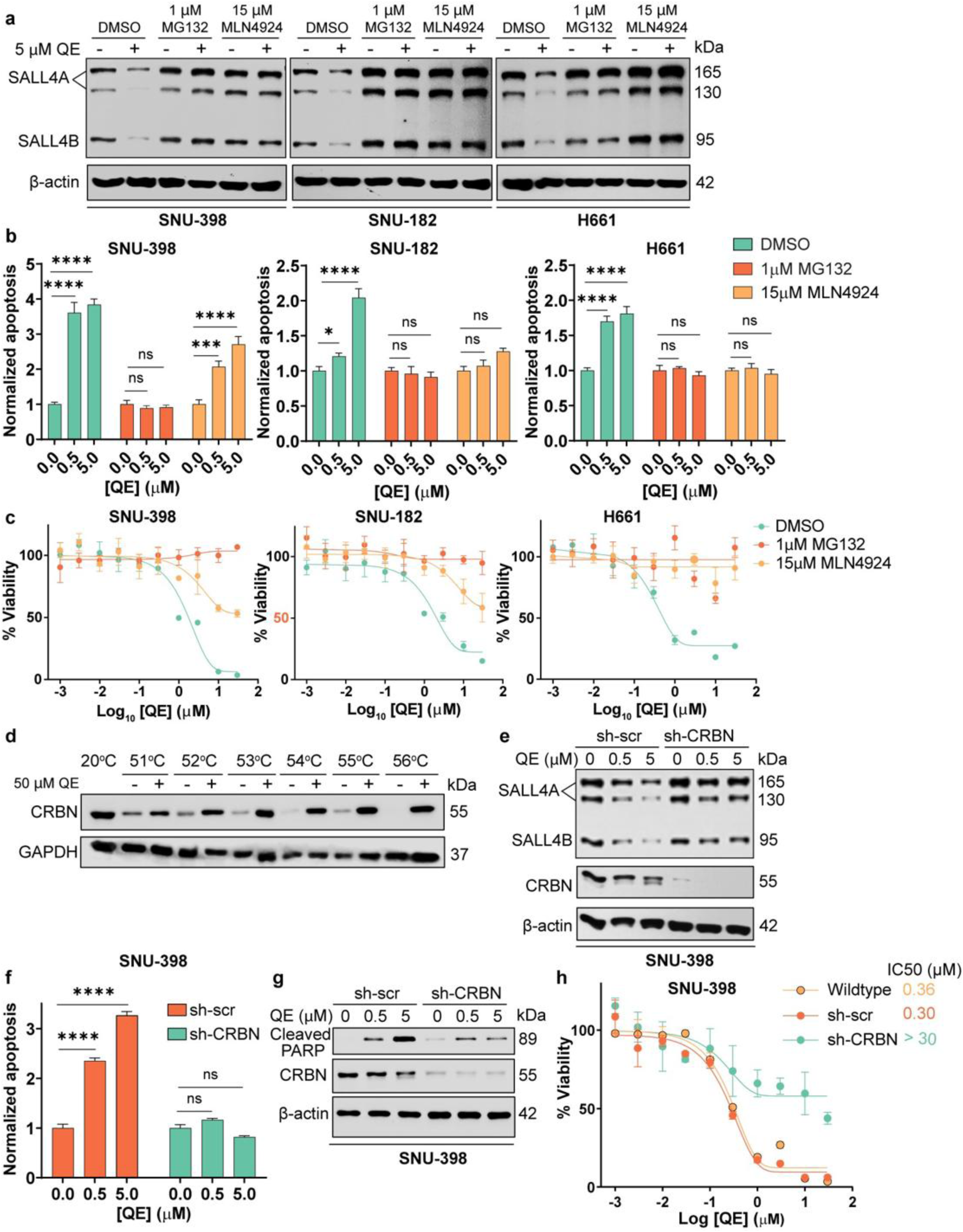
QE degrades SALL4B via CRBN, and CRBN disruption abrogates SALL4B degradation and downstream anti-cancer effects. **a,** QE-induced SALL4B degradation rescued by 18 hours co-treatment with MG132 (1 µM) or MLN4924 (15 µM) in SNU-398, SNU-182 and H661 cells. **b,** Caspase activity after 20 hours co-treatment with QE and MG132 or MLN4924 in SNU-398, SNU-182 and H661 cells (n = 3; mean ± SD). **c,** Viability of SNU-398, SNU-182 and H661 cells after 48 h co-treatment with QE and MG132 or MLN4924 (n = 3; mean ± SD). **d,** Immunoblot showing CRBN thermal stabilization after 4 h incubation of SNU-398 lysates with QE (50 µM). **e,** Immunoblot showing reduced SALL4B degradation in CRBN-deficient SNU-398 cells treated with QE for 18 h (representative of n = 3). **f,** Caspase activity in SNU-398 cells with normal or deficient CRBN levels after 24 hours QE treatment (n = 3; mean ± SD)**. g,** Immunoblot showing rescue of PARP cleavage in CRBN-deficient SNU-398 cells after 24 hours QE treatment (representative of n = 3). **h,** Viability of SNU-398 cells after 72 hours QE treatment with or without CRBN knockdown (n = 3; mean ± SD). **Statistics:** For (b) and (c), one-way ANOVA was performed separately for each condition (QE with DMSO, 1 µM MG132 or 15 µM MLN4924), followed by Dunnett’s multiple comparisons test versus the 0 µM control. For (f), one-way ANOVA was performed separately for each condition (sh-Scr or sh-CRBN), followed by Dunnett’s multiple comparisons test versus the 0 µM control. *P < 0.05, **P < 0.01, ***P < 0.001, ****P < 0.0001; ns, not significant.

To identify the E3 ligase responsible for QE-driven SALL4 degradation, we performed Cellular Thermal Shift Assay (CETSA). Thermal profiling of CRBN in SNU-398 cell lysates revealed pronounced stabilization upon QE treatment, with a thermal shift of up to 5°C (51 to 56°C), indicating direct engagement of cereblon (Fig. 5d).

We next assessed the functional requirement of CRBN. Genetic knockdown of CRBN in SNU-398 cells prevented QE-induced SALL4 degradation (Fig. 5e) and reduced QE-triggered apoptosis, as evidenced by diminished cleaved PARP levels and decreased apoptotic activity (Fig. 5f, g). Consistently, CRBN-knockdown and knockout SNU-398 cells showed increased viability following QE treatment compared to wild-type cells, whereas the viability of SALL4-independent SNU-387 cells remained unaffected by CRBN-knockout upon QE exposure (Fig. 5h, Supplementary Fig. 5b).

Together, these data demonstrate that a CRBN-dependent, proteasomal degradation of SALL4 is induced by QE, and provide a mechanistic insight into its mode of action.

### QE demonstrates potent anti-tumor effects across cell line-derived and patient-derived xenografts

We first evaluated the therapeutic potential of QE *in vivo* using a SALL4-high hepatocellular carcinoma (HCC) xenograft model derived from SNU-398 cells. In an initial tolerability and efficacy study, daily intraperitoneal administration of QE (10mg/kg) for 10 days significantly slowed tumor progression, resulting in a 64% reduction in tumor size relative to controls (Fig. 6a, b). Treatment was well tolerated, with no significant changes in animals’ body weight or spleen weight (Supplementary Fig. 6a, b).

**Figure 6.**
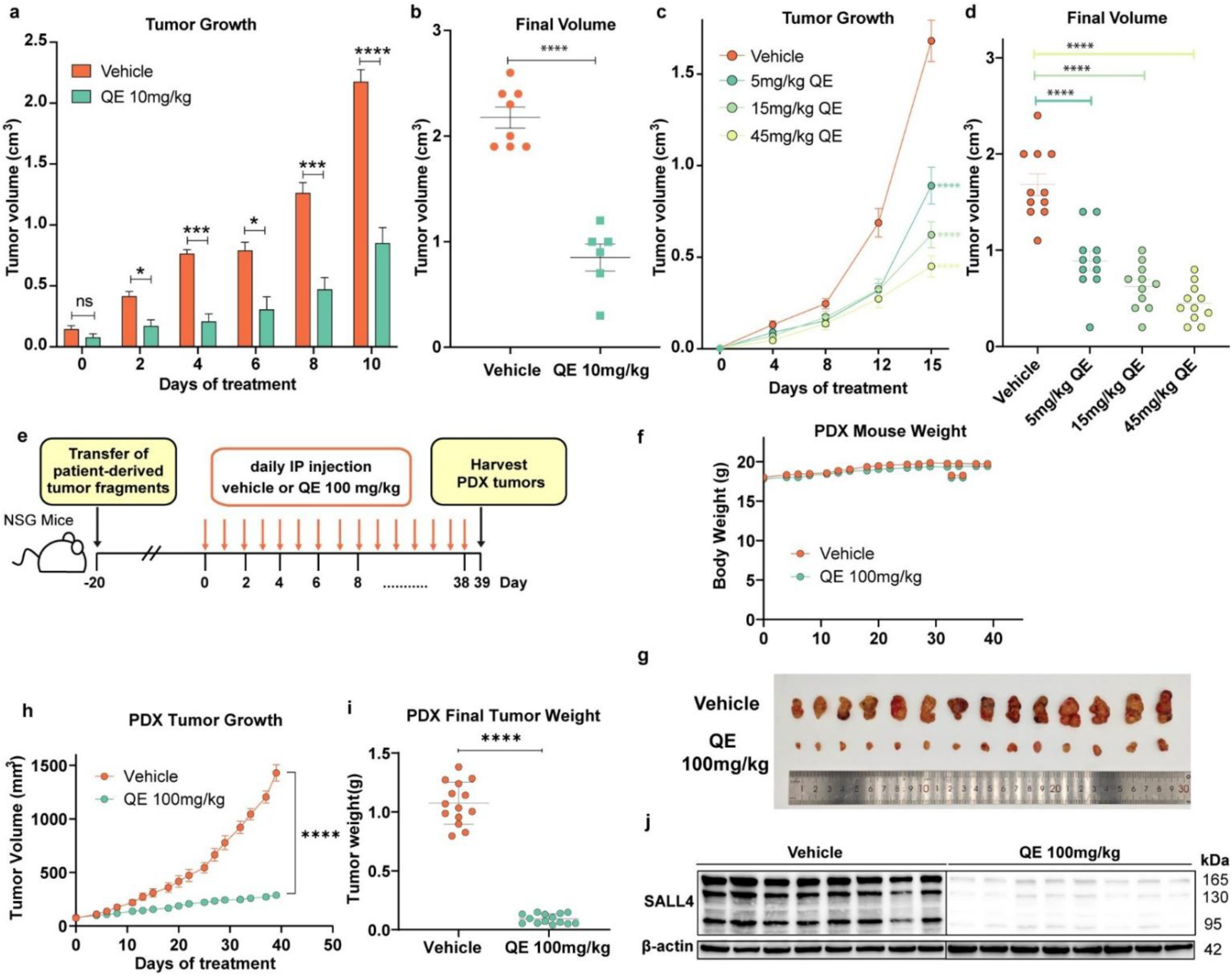
*In vivo* anti-tumor effect of QE. **a,** Tumour volume in mice treated with vehicle or QE (10 mg/kg) over 10 d (vehicle, n = 8; QE, n = 6; mean ± SEM). b, Final tumour volumes after 10 d of treatment (mean ± SEM). **c,** Tumour volume in mice treated with vehicle or QE at 5, 15 or 45 mg/kg for 15 d (n = 11 per group; mean ± SEM). **d,** Final tumour volumes after 15 d of treatment (mean ± SEM). **e,** Schematic of QE dosing regimen for dose response study**. f,** Body weight in vehicle- and QE-treated mice from day 0 to day 39 (n = 14; mean ± SEM). **g,** Representative tumour images from vehicle- and QE-treated mice (n = 14), with typical tumours from each group at end of treatment (n = 10). **h,** Tumour volume in vehicle- and QE-treated mice from day 0 to day 39 (n = 14; mean ± SEM). **i,** Final tumour weight on day 39**. j,** Immunoblot of SALL4 protein in tumour samples collected at day 39 (n = 8). **Statistics:** Two-way repeated-measures ANOVA with Dunnett’s multiple comparisons test for tumour volume studies in (a), (c) and (h), comparing each treatment group to vehicle at each time point. Unpaired Welch’s t-test for (b) and (i). One-way ANOVA with Dunnett’s post hoc test vs vehicle for (d). *P < 0.05, **P < 0.01, ***P < 0.001, ****P < 0.0001; ns, not significant.

To define the therapeutic window, we next tested daily QE administration at 5, 15, and 45 mg/kg for 2 weeks in mice bearing SNU-398 xenografts (Supplementary Fig. 6c). QE suppressed tumor growth at all doses tested (Fig. 6c), with dose-dependent reductions in tumor weight of 40%, 60%, and 70%, respectively (Fig. 6d).

Notably, even the highest dose was well tolerated, with negligible weight loss and normal spleen and liver weights (Supplementary Fig. 6d-f). These data establish the *in vivo* efficacy and tolerability of QE in a cell line-derived xenograft model of SALL4-high HCC.

Because patient-derived xenografts (PDX) more faithfully recapitulate human tumor biology and better predict clinical outcomes of experimental therapeutic agents than cell line-derived xenografts(17), we next used a SALL4-high HCC PDX model to evaluate whether QE could inhibit the growth of patient-derived tumors (Fig. 6e). Daily intraperitoneal QE treatment (100mg/kg) for 38 days was well tolerated with stable body weight (Fig. 6f). QE significantly inhibited tumor growth, achieving a mean tumor growth inhibition (TGI) of 79.8% (Fig. 6g-i). Antitumor efficacy was accompanied by SALL4 degradation in treated tumor (Fig. 6j).

Importantly, QE treatment did not alter liver or renal function, haematological parameters, or the morphology or weights of major organs (Supplementary Fig. 7a-d).

Together, these results demonstrate that QE exerts potent and well-tolerated antitumor activity in both cell line-derived and patient-derived xenograft models, supporting its promise as a therapeutic agent for SALL4-dependent cancers.

### SALL4B degradation impairs DNA damage response and replication, leading to loss of cancer cell viability

To understand the transcriptional programs governed by SALL4B that underlie its role in cancer cell survival, we performed RNA sequencing (RNA-seq) following SALL4B-specific knockdown in SNU-398 cells (Supplementary Fig. 8a). Differential expression analysis identified 2784 genes with FDR < 0.05 and log_2_FC > 0.5. Pathway enrichment using the Reactome database(18) revealed 23 significantly dysregulated pathways, including histone deacetylation, DNA methylation, β-catenin-TCF signaling, Notch signaling, DNA damage response, telomere packaging, and p53-regulated cell-cycle arrest (Supplementary Fig. 8b).

To delineate downstream pathways perturbed by QE-mediated SALL4B degradation, we compared transcriptomes from SALL4B-knockdown and QE-treated SNU-398 cells (Supplementary Fig. 8c, Supplementary Table 4). Of the 2784 DEGs from SALL4B knockdown and 2245 DEGs from QE treatment (FDR < 0.05 and log_2_FC > 0.5), 436 genes were shared (Fig. 7a, Supplementary Table 5). Reactome analysis of these common genes indicated prominent disruption of pathways involved in DNA replication/S-phase progression and DNA damage response (Fig. 7b).

**Figure 7.**
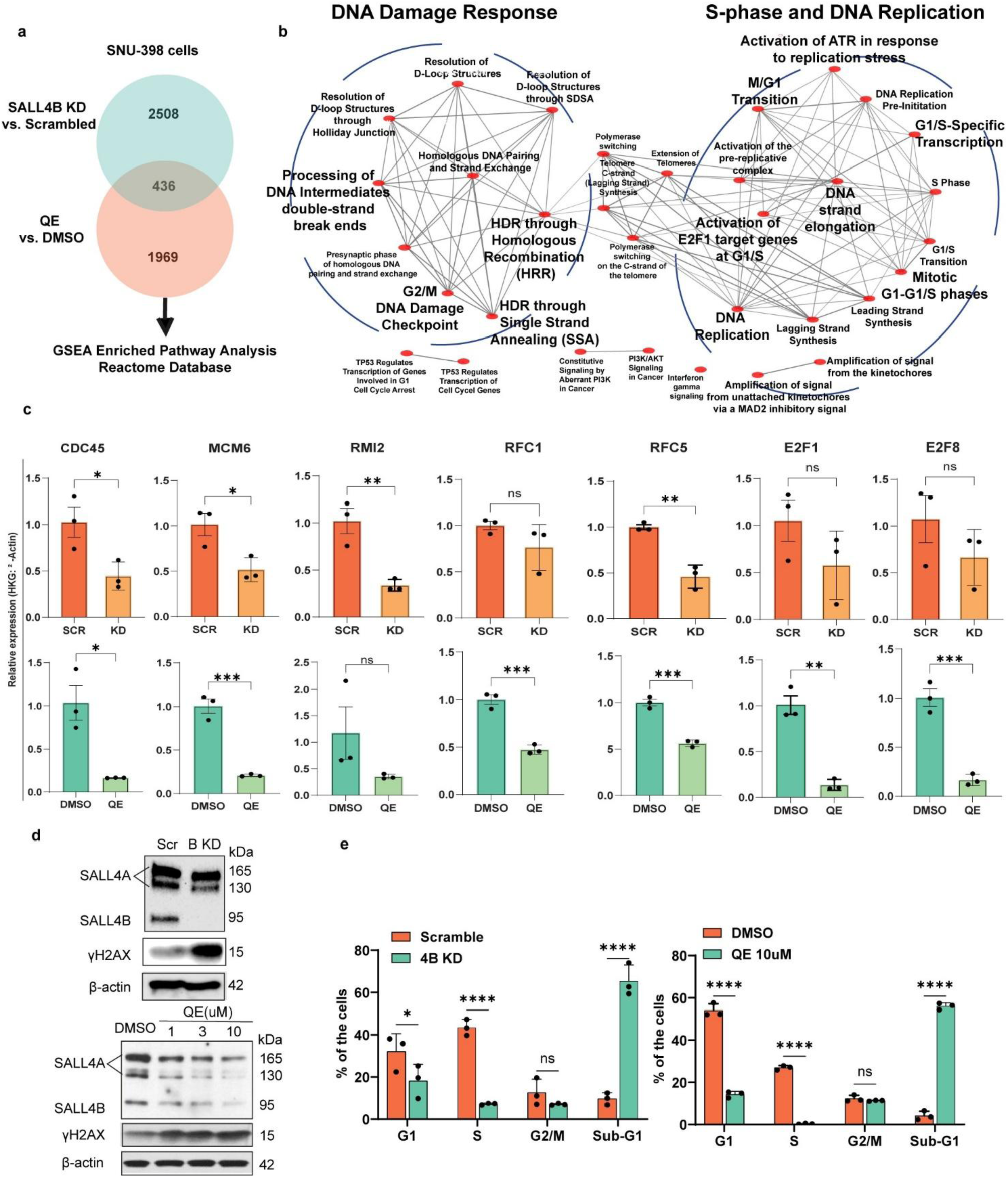
SALL4B degradation downregulates DNA damage response and replication in HCC. **a,** Venn diagram showing the number of differentially expressed genes (DEGs) with FDR < 0.05 and log2FC > 0.5 from RNA-seq analysis, overlapped or distinct upon SALL4B knockdown or QE treatment in SNU-398 cells. **b,** Pathway enrichment analysis of overlapping DEGs from SALL4B knockdown or QE treatment, highlighting DNA damage response and cell cycle pathway downregulation. **c,** qPCR-validated decrement in mRNA expression of selected genes in the DNA damage and cell-cycle-related pathways 48 hours post-SALL4B knockdown and 18 hours after 5 μM QE treatment in SNU-398 cells (n = 3; mean). **d,** Immunoblot for γ-H2AX DNA damage marker shows upregulation upon 7 days post SALL4B knockdown or 72 hours after QE treatment (representative of n = 2). **e,** Edu incorporation and cell-cycle distribution in SNU-398 cells after SALL4B knockdown or QE treatment (n = 3; mean ± SD), confirming the impairment of S-phase. **Statistics:** Unpaired two-tailed t-tests for qPCR validation in (b). Two-way ANOVA with Šídák’s multiple comparisons test for (e) comparing Scr vs KD and DMSO vs QE within each cell-cycle phase. *P < 0.05, **P < 0.01, ***P < 0.001, ****P < 0.0001; ns, not significant.

We validated key genes within these pathways by qPCR, including CDC45, MCM6, RMI2, RFC5, RFC1, E2F1, and E2F8, all of which were downregulated following SALL4B depletion by either QE or shRNA, mirroring the expression decreases observed in the RNA-seq data (Fig. 7c). some of these genes (CDC45, RFC5, MCM6) contribute to ATR activation in response to replication stress, suggesting compromised checkpoint function.

As a result of blockade in DNA Damage response and replication, we expect that γH2AX level increases(19). γH2AX evaluation demonstrated a substantial increase following QE treatment or SALL4B knockdown (Fig. 7d). Moreover, EdU incorporation assay demonstrated a marked reduction in S-phase cells and accumulation of ∼60% of cells in sub-G1/dead fractions in both conditions (Fig. 7e), indicating impaired DNA replication and cell-cycle progression.

Collectively, these transcriptomic and functional analyses show that SALL4B inhibition, either through QE-mediated degradation or genetic knockdown, leads to the coordinated loss of DNA replication and DNA damage response gene expression, ultimately impairing cancer cell viability (Fig. 8).

**Figure 8.**
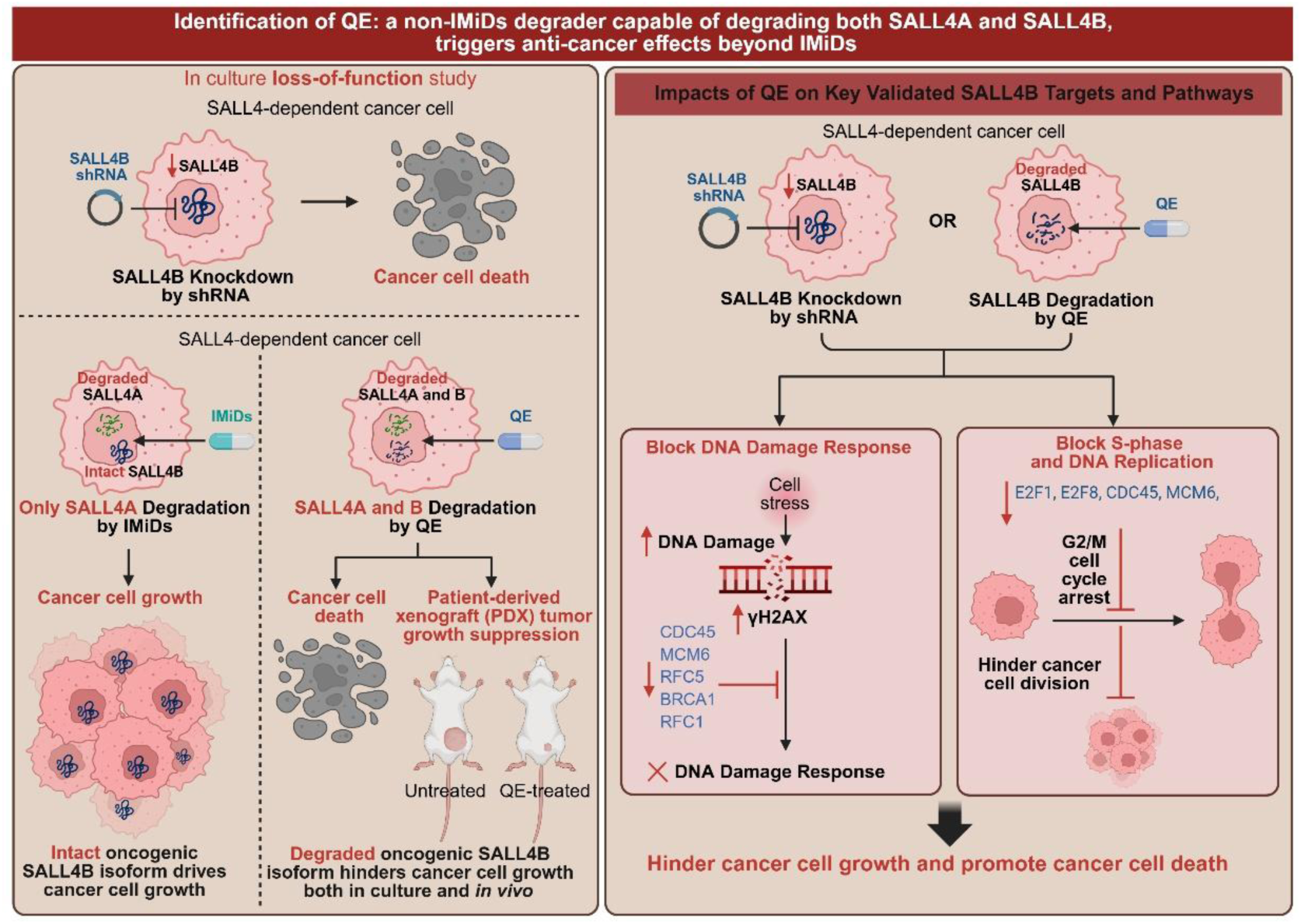
Schematic model depicting importance of SALL4B for cancer cell survival and targeting it by QE as a strategy to target the cell cycle and DNA-Damage pathways and killing SALL4-positive cancer cells.

## Discussion

Motivated by earlier studies that established SALL4 dependency in many solid tumors and leukemia(4, 7–10, 12, 13), our work sought to develop a small molecule SALL4 degrader as therapy for SALL4-mediated cancers. While developing our compound, two groups reported that immunomodulatory drugs (IMiDs) could degrade SALL4(14, 15). However, when tested across multiple SALL4-driven cancer cells, we found that IMiDs selectively degraded the SALL4A isoform while leaving SALL4B intact. This isoform selectivity is explained by the presence of IMiD-induced CRL4^CRBN^ binding motif (amino acids 410-433(14, 15)) within zinc finger cluster 2 (ZFC2) in SALL4A but not SALL4B. Importantly, IMiD-mediated SALL4A degradation failed to decrease the survival of SALL4-dependent cancer cells. Our *in vitro* SALL4B-knockdown studies, combined with the *in vivo* tumorigenic activity of SALL4B and prior evidence of SALL4B-driven leukemogenesis(16), support that effective therapeutic targeting of SALL4 requires degradation of the SALL4B isoform.

To address this unmet need, we developed an isoform-focused high-throughput degrader screen and identified QE as a small-molecule chemical probe capable of degrading SALL4B. QE induced potent, proteasome-dependent degradation of SALL4B and SALL4A across HCC and lung cancer cell lines and required CRBN for its activity. Furthermore, QE’s anti-cancer effects were dependent on SALL4 depletion, especially SALL4B, and QE produced pronounced anti-tumor activity in both cell line-derived and patient-derived HCC xenograft models without detectable toxicity. These results identify QE, rather than IMiDs, as a mechanistically informative chemical probe that provides a foundation for future therapeutic development targeting SALL4-mediated cancers.

The transcriptional regulatory functions of oncogenic SALL4B have not been described before. Our transcriptomic analyses revealed that SALL4B controls key genes involved in DNA replication, S-phase progression, and the DNA damage response. Among downregulated genes, CDC45, RFC5, and MCM6, components required for ATR activation during replication stress, are strongly associated with HCC progression(20–22). Additional genes within the enriched “homology-directed repair” pathway (BRCA1, RFC1, RFC5, RMI2) also exhibited reduced expression. RFC family members serve as prognostic markers in diverse cancers and function in DNA replication, repair, and cell-cycle control(23–26), while RMI2 is a prognostic marker for HCC(27–29). Functional assays confirmed that SALL4B inhibition induces DNA damage (γH2AX accumulation), disrupts S-phase entry, and leads to extensive cell death. Together, these findings suggest that SALL4B loss destabilizes DNA replication and repair programs, providing a mechanistic basis for the cytotoxicity observed in SALL4-dependent cancer cells following QE treatment.

Specifically, we note that IMiDs are known to degrade SALL4A, which plays a role in embryonic development; the potential teratogenicity of dual SALL4A/B degraders warrants careful evaluation. Since QE induces degradation of both isoforms, embryonic toxicity remains a theoretical concern, and future work is needed to evaluate its impact on pregnancy and development.

Against this backdrop, an important insight emerging from this study is that, although immunomodulatory drugs (IMiDs) have been highly effective in multiple myeloma and other hematologic malignancies (30), their intrinsic selectivity for SALL4A limits their therapeutic applicability in solid tumors, where SALL4B plays a critical role in sustaining cancer cell survival. This distinction has important implications for the field, which has largely avoided targeting SALL4 because of its historical association with thalidomide-induced teratogenicity (31). Our findings suggest that rationally designed small-molecule degraders capable of engaging SALL4B while bypassing canonical IMiD mechanisms may overcome this long-standing barrier and open a new therapeutic opportunity in SALL4-positive solid tumors. Building on these results, we are actively developing SALL4B-selective degraders to further minimize potential SALL4-associated effects on development.

Overall, using SALL4-mediated cancer as a model, our work highlights the need to understand how distinct isoforms contribute to tumorigenesis, treatment resistance, and therapeutic response. We introduce an isoform-focused screening platform that enables the rapid identification of small molecules capable of eliminating pathogenic protein isoforms. This approach provides a conceptual and technical framework for developing next-generation therapeutics targeting disease-driving protein isoforms across a broad spectrum of disorders.

## Materials and Methods

### Sex as a biological variable

The NSG mice that were used for Cell Line-Derived and Patient-Derived Xenografts were both male and females and and similar findings are reported for both sexes.

### Antibodies

Western blot primary antibodies used were SALL4 EE30 (sc-101147), Beta-actin-HRP (sc-47778 HRP), GAPDH-HRP (sc-47724 HRP) from Santa Cruz Biotechnology; C-myc (D84C12), Cleaved-PARP (D214), and CRBN (NBP1-91810) from Novus Bio. Secondary antibodies were goat anti-mouse IgG-HRP (sc-2005 or sc-516102) and goat anti-rabbit IgG-HRP (sc-2004 or sc-2357) from Santa Cruz Biotechnology.

### Cell Culture

All cell lines were obtained from ATCC. Human hepatocellular carcinoma (HCC) cell lines SNU-398, SNU-182, SNU-387, SNU-449, SNU-475, and non-small-cell lung carcinoma (NSCLC) cell lines H661 and H1299 were cultured in filter-sterilized RPMI medium (Gibco) with 10% FBS (Hyclone GE Healthcare), and 1% penicillin-streptomycin (Gibco) at 37°C, 5% CO_2_. Cells were checked and confirmed mycoplasma-negative using a MycoAlert detection kit (Lonza).

CAL51 (CVCL_1110) was obtained from DSMZ (Leibniz Institute DSMZ-German Collection of Microorganisms and Cell Cultures GmbH) and cultured according to the provider’s instructions in the absence of antibiotics. Cells were maintained in Gibco™ DMEM, High Glucose (Thermo Fisher Scientific, Cat# 11965118) supplemented with 10% fetal bovine serum (Gemini Bio).

### Drug Treatment

The drug library used in this study was from Novartis Institutes for Biomedical Research (NIBR), Cambridge, and consisted of 50,000 small molecules(32, 33) (> 95% purity). The IMiDs used for comparison studies were Pomalidomide (>98% purity, Sigma-Aldrich), Lenalidomide (99.82% purity, Selleckchem, TX, USA), Thalidomide (>98% purity, Sigma-Aldrich), CC-122 (99.73% purity, Selleckchem, TX, USA), CC-220 (98.67% purity, Selleckchem, TX, USA), and CC-885 (99.23%, MedChemExpress, NJ, USA). MG132 (>97% purity), and pevonedistat MLN4924 (99.01% purity) were from Selleckchem, TX, USA.

The QE compound(34), namely 1-(2-hydroxy-4-nitrophenyl)-3-(2-methoxyphenyl) urea (Pubchem 9796445) used has a > 95% purity from NIBR, Cambridge, USA, and > 98% purity from Enamine. The purity was assessed by NMR and/or HPLC.

### Production of SALL4 (1-300aa) Protein in E.coli for High throughput- Screening

DH5alpha were transformed with pET His6 TEV LIC Cloning vector (2B-T) containing SALL4 (1-300aa). pET His6 TEV LIC cloning vector (2B-T) was a gift from Scott Gradia (Addgene plasmid # 29666).

SALL4(1-300aa) cloned into this vector was expressed with the N-terminal 6-His tags in the E. coli BL21(DE3)/pLysS strain using a T7 expression system. SALL4 1-300 was purified by His trap Ni^2+^ column and gel filtration. After purification, the SALL4(1-300) protein was further characterized by SDS-PAGE Gel and mass spectroscopy for purity and check of identity. A fluorescent polarity assay was used to confirm the ability of the produced SALL4 (1-300aa) to form a complex with its RBBP4 protein partner. Biophysical assays, circular dichroism, and a thermal shift assay were used to evaluate the protein secondary structure and thermal stability, respectively. Dynamic Light Scattering (DLS) was used to detect and monitor any sign of protein aggregation. The yield of protein obtained from 1L of E.coli culture was 6mg. A total of 686mg of SALL4 1-300 protein was produced for drug screening and characterization assays.

### Thermal Shift Assay for High-throughput Screening (HTS) – Primary Screen

The library of 50,000 small molecules from the Novartis Biomedical Research Institute (NIBR)^29,30^ was screened at 50 μM and 100 μM against 13.8 μM of purified SALL4 (1-300aa) protein in a buffer containing 100mM potassium phosphate, 100mM NaCl at pH 7. The screening assay was performed in 384-well PCR plates using a LightCyler^®^ 480 Real-Time PCR System (Roche Applied Sciences). A 7-min run started at 30°*_C_* and ended at 90°*_C_*. Sypro Orange (Sigma-Alrich) was used as the sensitive fluorescent dye for monitoring thermal protein denaturation. Hits were defined as compounds that can induce a change in protein melting points 3 times higher than the standard deviations of the apo-protein melting point from the DMSO control.

### Dual-Luciferase Reporter Assay for High-throughput Screening (HTS) – Secondary Screen

The dual luciferase assay was performed in a 1,536-well plate that allows high throughput screening (HTS) of thermal shift hits from the primary screen, searching for SALL4 small molecule binders that can degrade the SALL4B isoform. The assay employed in this study was modified from a previously reported dual luciferase assay.(35) The SALL4A or B isoform was subcloned as nano-luciferase fusion (Nluc) into the Nluc/Fluc plasmid (pLL3.7-EF1a-IRES-Gateway-nluc-2xHA-IRES2-fluc-hCL1-P2A-Puro), which was provided by William Kaelin. H1299 cells that stably express SALL4A/B_Nluc_Fluc were seeded in a 1,536-well transparent bottom white plates (Corning) at 2,000 cells per well in 6 μL of RPMI medium and grown overnight (16h) at 37°C, 5% CO_2_. Cells were treated with 8 *_μM_* of compounds for 6h. The Nano-Glo® Dual-Luciferase® Reporter Assay System (Promega, Cat#: N1650) and a Pherastar plate reader were used for measuring the luciferase signal, following the manufacturer’s instructions. For data analysis, the Nanoluc luminescence was normalized to firefly luciferase reading and then compared to the DMSO control. The screen was performed in triplicates.

A counter-screen was also performed to eliminate false positive hits that interfere with the luciferase signal. The protocol for the counter screen is similar to that of the main screen, except that there was no incubation period of 6h. Hits were selected as compounds resulting in 50% or more reduction in Nluc/Fluc signal in the main screen with minimal interference (<15% signal reduction) of luciferase signal itself in the counter screen.

### SALL4(1-300aa) protein expression and purification

SALL4(1-300) was expressed using a His6-tagged construct a pET28a(+) expression vector and the *E. coli* BL21 (DE3) expression strain (New England Biolabs). Modified M9 minimal media for expression of U-^15^N-labeled SALL4(1-300) was prepared as follows: 6 g Na_2_HPO_4_, 3 g KH_2_PO_4_ and 0.5 g NaCl were dissolved in 1 L of double deionized water and autoclaved. Then, 35 mg Kanamycin was added, as well as sterile filtered solutions of MgSO_4_ (final concentration = 2.5 mM), CaCl_2_ (final concentration = 0.1 mM), 4 g D-Glucose, 1 g ^15^NH_4_Cl and 100 µL of a 10,000 x vitamin mix (5 g/L riboflavin, 5 g/L niacinamide, 5 g/L pyridoxine monohydrate and 5 g/L thiamine dissolved in ethanol). A culture in 10 mL LB per liter large culture was grown at 37 °C overnight (∼ 10 h) and used to inoculate 1L of the M9 media described above. The culture was grown in a shaking incubator at 37 °C until an OD_600_ of ∼ 0.4, after which the temperature was reduced to 20 °C. At an OD_600_ of 0.7-0.9, IPTG was added to a final concentration of 0.5 mM and the protein was expressed at 20 °C overnight. The next day, the bacteria were harvested by centrifugation at 6,000 x g for 15 min and the cell pellets were either stored at −80 °C or immediately prepared for purification.

Bacterial cell pellets were resuspended in ∼ 30 mL resuspension buffer (50 mM Tris-HCl, pH = 8.0, 350 mM NaCl, 10 mM imidazole, 2 mM β-mercaptoethanole [BME], 1 cOmplete™ Protease Inhibitor Cocktail tablet) per liter culture. Resuspended cells were then lysed by sonication (30 % amplitude, 5 min sonication time, 2 s on pulse, 4 s off pulse). Cell lysates were pelleted by centrifugation at 30,000 x g for 40 min at 4 °C. Cleared lysates were incubated with ∼ 3 mL Ni-NTA agarose resin, preequilibrated with wash buffer (50 mM Tris-HCl, pH = 8.0, 350 mM NaCl, 10 mM imidazole, 2 mM BME) per liter culture for 2 h at 4 °C. The resin was washed with wash buffer until 10 µL of eluent no longer stained blue with 50 µL of Bradford reagent (Coomassie blue G-250). Bound protein was eluted with elution buffer (50 mM Tris-HCl, pH = 8.0, 350 mM NaCl, 350 mM imidazole, 2 mM BME). Eluted proteins were dialyzed against 2 L dialysis buffer (20 mM sodium phosphate, pH = 6.8, 150 mM NaCl, 2 mM DTT, 1 mM EDTA) after addition of TEV protease (∼0.2 g purified TEV per liter culture) to cleave His6-tags for 16 h at 4 °C. Cleaved protein was concentrated to ∼ 5 mL and subjected to size exclusion chromatography using a Superdex 75 increase 10/300 GL column preequilibrated with SEC buffer (50 mM sodium phosphate, pH = 6.8, 150 mM NaCl, 2 mM DTT, 1 mM EDTA).

### Cell Titre Glo Viability Assay

The assay was performed in a 384-well plate (Corning 3570). SNU-398 and SNU-182 cells were seeded at 1,500 cells per well in 40 μL of RPMI medium. For SNU-387, SNU-449, SNU-475, the number of cells per well was 750. For H661 and H1299, there were 1,000 cells per well. Cells were then incubated overnight at 37°C, 5% CO_2_ and treated with compounds or DMSO vehicle at desired concentrations for 72 or 96 hours. The CellTiter-Glo Assay System (Promega, Cat#: G7573) and Pherastar or Tecan Infinite M200 Pro plate reader were used according to manufacturer’s instruction for measuring the luminescence.

### Colony Formation Assay

SNU398, SNU-387, SNU-182, SNU-449, SNU-475, H66,1 and H1299 cells were trypsinized and counted 72 hours after transduction. 5,000-10,000 cells were seeded per well in a 6-well plate and incubated at 37°C, 5% CO_2_ for 7 days. The cells were fixed with 10% neutral buffered formalin solution (Sigma-Aldrich) and stained using Crystal Violet (Sigma-Aldrich). Stained wells were washed with water until the excess dye was removed and left to dry overnight. The colonies were imaged and quantified using ImageJ.

### Soft Agar Assay

A base agar layer (0.6%) was prepared by diluting 1.2% agar with 2X complete culture media and allowed to set in a 6-well plate. The cells were trypsinized post-treatment, counted, and diluted to a final concentration of 10,000 cells/well. A top agar layer (0.3%) containing single-cell suspension was made by diluting 0.6% agar with the cell suspension made in the previous step. The cells were incubated at 37°C, 5% CO_2_. The culture medium was changed every two days. Colonies were imaged and quantified using ImageJ.

### Caspase Glo Assay for Apoptotic Cell Analysis

The assay was performed in 384-well plates (Corning 3570). Cells were seeded at 5,000 cells per well in 30 μL of RPMI medium and incubated overnight at 37°C, 5% CO_2_. 10 μL of compounds or DMSO vehicle in RPMI (0.5% final DMSO) was added. After 20h or 24h of incubation, 30 μL of Caspase-glo reagent were added and the plate was allowed to sit at room temperature in the dark for 2 hours. The luminescence was monitored and recorded using a Tecan Infinite M200 Pro plate reader.

### SALL4 Knockdown by Lentiviral Transduction

shRNA targeting total SALL4 (sh-SALL4_1 & sh-SALL4_2)(4), SALL4B (sh-SALL4B) and a scrambled control (sh-scr)(4) with the following sequences were cloned into the pLL.3 vector containing GFP:

sh-scr: 5’-GGTACGGTCAGGCAGCTTCT-3’

sh-SALL4_1: 5’-GCTATTTAGCCAAAGGCAAA-3’

sh-SALL4_2: 5’-GCGTTGAAACAGGCCAAGCTG-3’

sh-SALL4B: 5’-CACAAGTGTCGGAGCAGTC-3’

For lentiviral transduction, SNU-398 cells were spun with lentiviral particles containing the specified vectors in RPMI (plus 10% FBS, and 5μg/ml polybrene) at 800g for 60 minutes at 32°C. Transduction efficiency was assessed via GFP signal using a BD FACSAria™ (BD Biosciences). The cells were then incubated at 37°C, 5% CO_2_.

### Lentiviral CRBN Knockdown

shRNA targeting CRBN (sh-CRBN) or a scrambled control (sh-scr) with the following sequences were cloned into the pLKO.1 vector:

sh-scr: 5’-CAACAAGATGAAGAGCACCAA-3’

sh-CRBN: 5’-CAGGATAGTAAAGAAGCCAAA-3’

For lentiviral transduction, SNU-398 cells were spun with lentiviral particles containing the aforementioned vectors in RPMI (plus 10% FBS, 8μg/ml polybrene) at 800g for 60 minutes at room temperature. 96h post-infection, cells were seeded in 6-well or 384-well plates and incubated overnight at 37°C, 5% CO_2_. Cells were then treated with DMSO or QE at desired concentrations (0.5% final DMSO) for 18h and 24h.

Immunoblot analysis was subsequently performed with the indicated antibodies. For cell viability, cells were treated for 72h at 37°C, 5% CO_2_.

### CRISPR-mediated CRBN Knockout

The CAS9 vector from Addgene (Addgene plasmid # 110837) was used to produce lentiviral particles. SNU-398 and SNU-387 cells were then transduced with the virus in the presence of 8μg/ml polybrene, followed by selection with 2ug/ml of Puromycin for seven days.

An sgRNA targeting CRBN was designed using CRISPick gRNA design tool with the following sequence:

5’-ATAGTACCTAGGTGCTGATA-3’

The sgRNA was cloned into the FgH1tUTG vector (Addgene plasmid #70183) and the CAS9-positive SNU-398 and SNU-387cells were spun with the viral particles and were selected based on the GFP selection.

### Lentiviral SALL4A or B Isoform expression

Lentiviral constructs for SALL4A and SALL4B cloned in pFUW were used for expression. For lentiviral transduction, SNU-398 cells were spun with viral particles containing empty pFUW vector, SALL4A or SALL4B in RPMI (plus 10% FBS, and 5μg/ml polybrene) at 800g for 60 minutes at 32°C. Transduction efficiency was assessed via mCherry signal using a BD FACSAria™ (BD Biosciences). Following 3 days post transduction, the cells were selected using puromycin (1μg/ml) for one week. Expression was verified using qRT-PCR and western blot. Cells were subsequently seeded for analyzing drug sensitivity.

### Cellular Thermal Shift Assay (CETSA)

The CETSA assay was modified from Martinez-Molina et al.(36) SNU-398 cells were washed with cold PBS, harvested and centrifuged for 5 min at 1,200g. For cell lysis, cells were resuspended in 1X NP-40 lysis buffer containing 10mM Tris-HCl (pH 7.4), 150mM NaCl, 2.7mM KCl and 0.4% NP-40 and rotated end--over--end at 4°C for 30 mins. The lysate was centrifuged at 20,000 rpm for 10mins at 4°C and the supernatant was collected. Cell lysates were divided into two halves, with one being treated with QE and the other with DMSO control. After incubating at room temperature for the desired duration, the respective lysates were further subdivided into smaller aliquots of 40 μL and heated in a thermal cycler at different temperatures, ranging from 51°C to 56°C, for 3 minutes, followed by a 3-minute cool-down at room temperature. After centrifugation of the heated lysates at 15,000 rpm for 10 minutes at 4°C, the supernatants were collected and subjected to immunoblot analysis.

### SALL4B transgenic mice and two-stage chemical carcinogenesis

SALL4B transgenic mice were generated in a C57BL/6 background as previously described,(16) and were maintained at the mouse facility at Children’s Hospital Boston (CHB). All animal work was approved by the IACUC under protocol 10-10-1832. The SALL4B primer sequences for genotyping include the following: forward primer, 5’-AGCAGAGCTCGTTTAGTGAACCG-3’, and reverse primer, 5’-CTGTCATTCATGATGAGGACAGG-3’. For the two-stage chemical carcinogenesis experiment, both SALL4B transgenic and wild type male C57BL/6 mice received a single i.p. injection of 5 mg *N-*-nitrosodiethylamine (DEN)/kg body weight (Sigma, St. Louis, MO) in sterile water at 15 days of age. Mice were then administered 0.05% of phenobarbital (Sigma) in drinking water from 2 weeks after DEN injection (4 weeks of age). Mice were sacrificed 10 and 23 weeks after the start of the phenobarbital (PB) diet by carbon dioxide inhalation and necropsied. Deaths and moribund cases were also necropsied. Body weights were recorded, and livers removed, weighed and examined for grossly visible lesions. Each liver lobe was fixed in 10% neutral buffered formalin, trimmed, and embedded in paraffin. For routine histological analysis, two representative sections were prepared from each liver lobe, 4-6 µm sections were prepared from paraffin blocks and stained with H&E. Liver lesions were classified according to criteria defined previously(37, 38). The present of foci of cellular alteration was graded based on the percentage of foci area compared to the liver section as following: +: < 15%, ++: 15-40%, +++: 40-75%, ++++: 75-100%.

### Immunohistochemistry

Paraffin tissue sections of 4 µm were deparaffinized with Histoclear and hydrated in graded ethanol. Antigen retrieval was performed by heating in citrate buffer at 95°C for 30 minutes for Ki-67 IHC. Non-specific signals were blocked by peroxidase treatment for 10 minutes at room temperature, followed by protein block using Dako Protein Block Serum-Free reagent (Cat# X0909) for 30 minutes at room temperature. Primary antibodies were incubated at room temperature for one hour in a humidified chamber, followed by HRP-conjugated secondary antibody incubation for 30 minutes at room temperature. Antibody binding was revealed by DAB and reactions were stopped by immersion of tissue sections in distilled water once a brown color appeared.

Tissue sections were counterstained by hematoxylin, dehydrated in graded ethanol and mounted. The following antibody was used: Ki-67 (Novus Biologicals, Littleton, CO, USA #NB110-89717). All reagents for immunohistochemistry were from Dako (Dako, Glostrup, Denmark A/S). Appropriate positive and negative controls were included for each run of IHC.

### HCC Cell Line-Derived Xenografts

Animals were maintained, and all animal work was performed following the protocols approved by the Institutional Animal Care and Use Committee. The SNU-398 cell line was cultured as described above in ‘Cell Culture’. To establish the SNU-398 xenografts, NOD. Cg − Prkdc^scid^ Il2rg^tm1Wjl^/SzJ (NSG) mice (6-8 weeks) were anesthetized using 2% isoflurane USP (Baxter), and 0.5 × 10^6^SNU-398 cells in 200*_μL_* of 1:1 RPMI:Matrigel were subcutaneously injected to the mouse flanks. When tumors were palpable, mice were randomized to receive either QE treatment or vehicle containing 5% DMSO, 12.5% Ethanol, and 12.5% Cremophor EL (Sigma-Aldrich) in 70% 1X PBS. Freshly prepared vehicle or QE was administered by intraperitoneal (i.p.) injection, once daily (qd) at 10mg/kg in the pilot study and 5, 15, and 45mg/kg for dose-expansion studies. Before each drug injection, mouse weight and tumor volume were monitored. Mice were euthanized once tumors reached more than 1.5 cm in diameter. The tumors were harvested, imaged and snap-frozen for storage at −80°C until later use.

### HCC Patient-Derived Xenograft Model

Patient-derived xenograft (PDX) studies were conducted at the Institute of Model Animal, Wuhan University. All clinical sample acquisitions were approved by the Medical Ethics Committee, Zhongnan Hospital of Wuhan University (No. 2022076K), with informed consent obtained from the patient. Fresh liver cancer samples were obtained from patients undergoing surgery at Zhongnan Hospital of Wuhan University. All patients were pathologically confirmed to have hepatocellular carcinoma (HCC). After collection, tumor samples were washed 2-3 times in pre-cooled saline containing penicillin, kanamycin sulfate, and amphotericin B, then cut into small tumor pieces of 1-2 cm³. A portion of the tissue was reserved for Western blot and qPCR to assess SALL4 expression, while the remaining tissue was stored in pre-cooled DMEM medium for future use.

NSG mice, aged 4-6 weeks (purchased from Beijing HFK Bioscience Co., Ltd.), were selected as recipient mice. An incision was made at the base of the hindlimb of NSG mice, and a tumor block was inserted through the incision into the axillary region of the forelimb using a trocar. The trocar was then withdrawn, and the incision site was disinfected with povidone iodine-. Tumor growth was monitored, and when each transplanted tumor reached a size of 0.4-0.8 cm³, it was excised from the host and passage into new NSG mice. Ultimately, a third generation- PDX model derived from a 72-year-old male patient with hepatocellular carcinoma exhibiting high SALL4 expression was used for subsequent experiments.

When the average tumor volume in the PDX model reached 80 mm³, the mice were randomly divided into two groups: a vehicle control group and a QE treatment group receiving a dose of 100 mg/kg. Freshly prepared drugs or solvents were administered via intraperitoneal injection once daily. The health status of the mice was monitored before each administration, and body weight and tumor volume were measured at least three times per week. When the tumor diameter exceeded 1.5 cm³, blood samples were collected from the eyeballs of the mice, which were then sacrificed. Tumors were harvested, rapidly frozen, and stored at −80°C for further use.

### Transcriptomic Profiling

For knockdown, SNU-398 cells were seeded 0.1 × 10^6^ cells per well, transduced with scramble and SALL4B shRNA and collected at 48 hours. For drug treatment, SNU 398 -cells were seeded at 0.1 × 10^6^ cells per well, treated with QE at 5μM and collected at 18 hours. Subsequently RNA was purified employing the RNeasy Mini kit (Qiagen). The sample quality was confirmed with Bioanalyzer (Agilient, Santa Calra, CA), before sending for whole transcriptome sequencing using Hi-seq 2500 sequencer. All samples yielded an RNA integrity number of > 9. For data processing, raw fastq files first had optical duplicates removed using clumpify from the BBTools suite(39) using the flag “dedupe spany addcount”. Next, adaptor trimming was performed using bbduk, also from the BBTools suite using the flag “ktrim=l hdist=2 ref=<adapter_path>”. Next, reads were quality trimmed with Trimmomatic(40) in paired end mode with the flags “LEADING:28 TRAILING:28 SLIDINGWINDOW:4:26 MINLEN:20”. Alignment was performed using STAR v2.7.9a(41) with the flags “--outFilterScoreMinOverLread 0.25 --outFilterMatchNminOverLread 0.25 --outFilterMultimapNmax 5“ with the resultant bam files sorted and indexed using SAMtools(42). BamCoverage(43) was used to generate coverage maps using default parameters. Raw counts were generated using featureCounts(44) with flags “-M -O -J -f” using the Ensembl v101 annotation release for Homo Sapiens hg38. Differential gene expression analysis was performed using a custom R script based on an available tutorial (https://combine-australia.github.io/RNAseq-R/06-rnaseq-day1.html). Briefly, cpm counts were calculated and genes with cpm values < 0.5 were excluded. Normalization factors were then calculated for TMM normalization. Differential expression was calculated using limma-voom using appropriate design matrices.

Finally, appropriate contrasts were applied and differential gene expression tables generated for the appropriate test conditions. Differential gene expression lists were then uploaded to Qiagen’s Ingenuity Pathway Analysis suite(45) and analyzed online. Differentially expressed genes with FDR < 0.05 and log2FC (Fold Change) > 0.5 were used for pathway enrichment analysis by Reactome database in CluGo plugin (Version 1.5.5) of Cytoscape software (3.9.1). Sequencing data have been deposited in GEO with accession number GSE228864 (reviewer access token: wlijwiyclzyrroz).

### Quantitative RT-PCR

Extracted RNAs were reverse-transcribed using qScript cDNA Synthesis Kit (Quantabio). Exon-junction primers for CDC45, MCM6, RFC5, RFC1, RMI2, E2F1, and E2F8 were designed by AlleleID version 7.6 (Primer Biosoft, Palo Alto, USA). Primer specificity was validated by NCBI BLAST and the expression level of the genes was assessed by GoTaq® qPCR Master Mix (Promega), and QuantStudio 5 (Applied Biosystems). β-ACTIN was used as the internal control. The nonparametric Mann-Whitney test with a p-value < 0.05 was applied to show significant expression changes by GraphPad Prism 9 Software.

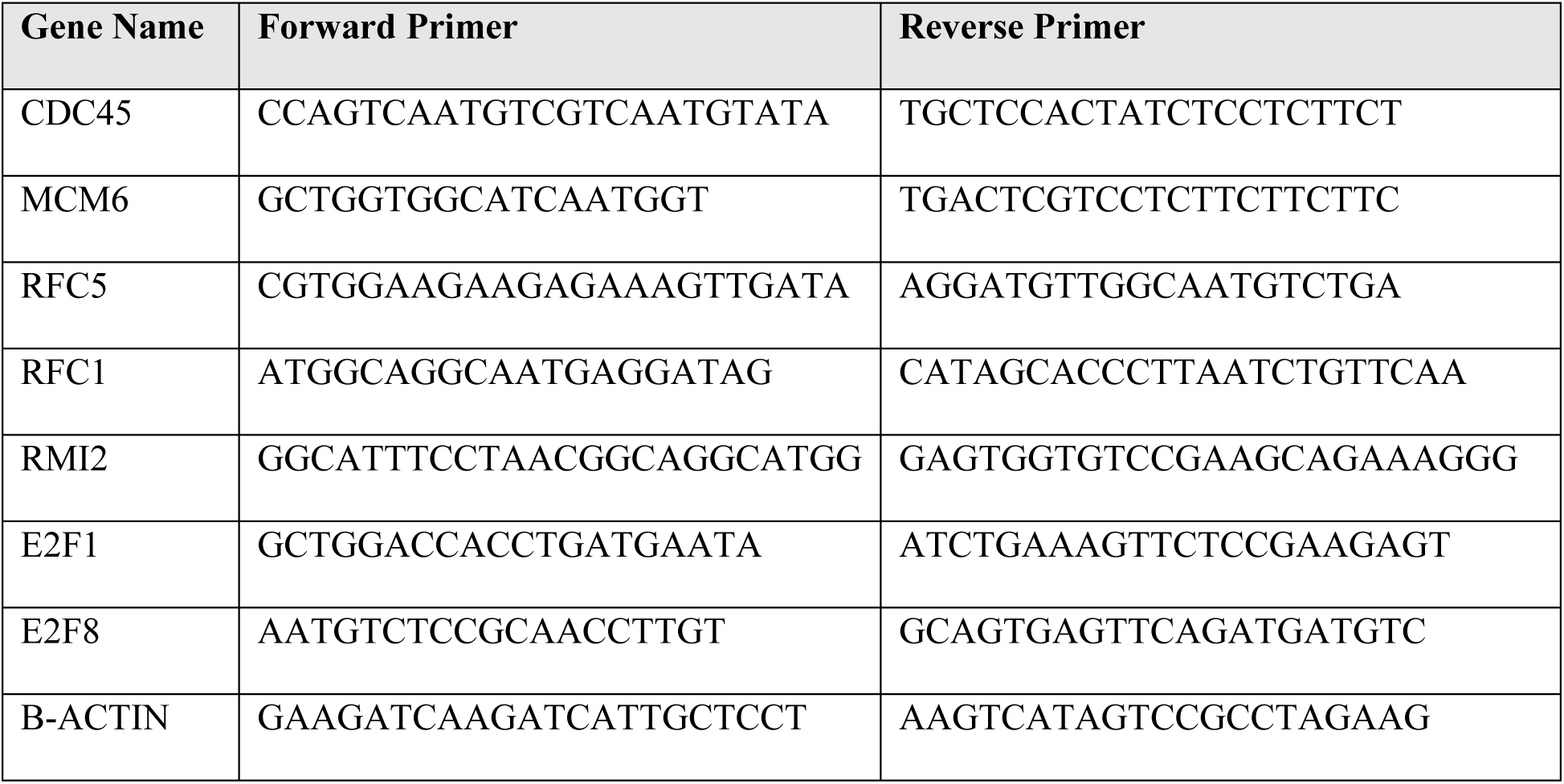

### Edu Incorporation assay

The Click-iT™ EdU Alexa Fluor™ 488 Flow Cytometry Assay Kit (ThermoFisher) was used for cell cycle analysis(C10420). The suspended cells were treated with Edu using a concentration of 10 μM for 5 hours. All the fixation, permeabilization and washing steps were done according to the manufacturer protocol except substituting Alexa Fluor® 488 azide with CalFluor 647 azide (Vector Laboratories) as the SALL4B KD and Scramble cells contained GFP marker. After staining the cells and washing with PBS/BSA 1%, the cells were treated with 2ug/ml of FxCycle™ Violet (ThermoFisher) for 30 mins at room temperature to stain DNA. The samples were analysed using BD CantoII or BD SymphonyA1 flow cytometry machines using 405 nm excitation and 450 nm FxCycle and an excitation around 650 nm and emission 665 nm for AlexaFluor647.

### Data Availability Statement

The dataset supporting the conclusions of this article is available in the Gene Expression Omnibus under accession number GSE228864. Immunoblots source data are shown in Supplementary Files-Uncropped blot images.

### Statistics

In this study, we implemented a range of both parametric and non-parametric tests aligned with each experimental design, and GraphPad Prism 9 was used for data analysis. All details of each statistical analysis, according to the graphs, are provided in the legend of each figure, along with the thresholds and levels of significance.

### Study approval

SALL4B transgenic mice were developed at the mouse facility at Children’s Hospital Boston (CHB). All animal work was approved by the IACUC under protocol 10-10-1832. Patient-derived xenograft (PDX) studies were conducted at the Institute of Model Animal, Wuhan University. All clinical sample acquisitions were approved by the Medical Ethics Committee, Zhongnan Hospital of Wuhan University (No. 2022076K), with informed consent obtained from each patient.

## Author contributions

K.A.V. designed and performed the drug screening and validation assays, biological testing of drugs in cells and mouse xenografts with the help of B.H.L. and S.M. K.A.V, J.P.T. purified recombinant SALL4(1-300aa) protein, J.P.T., J.L.T., D.E.C., D.S.A., D.M., Q.Z.; K.K. designed and carried out isoform function studies in cells with the help of F.L.; C.G., X.T.; M.L., M.Y.D., Z.L., and G.W designed and performed the SALL4B transgenic mice experiments and the xenotransplant models ; A.J.S performed the experiment testing thalidomide sensitivity of the SALL4 isoforms mutants with different zinc -finger -cluster (ZFC) deletion; J.Q. and L.H.S. performed the CRBN biochemical assays; K.K., M.A.B., S.M., V.I. performed and analyzed the RNA-seq; S.M. performed all post RNA-seq analysis and designed and implemented the genetic modification tests; D.S.A., H.A., and P.D.F, D.M., G.W., L.C., D.G.T. supervised the project; J.L.T., J.Q., H.A., C.B., and C.X.S., contributed intellectually to the manuscript writing; K.A.V., K.K., S.M., L.C., D.G.T. wrote the manuscript with the contribution of all the authors.

## Acknowledgements

We thank William Kaelin (Dana-Farber Cancer Institute) for kindly sharing the dual-luciferase plasmid. We thank the Fastlab Program at the Novartis Institute for BioMedical Research (NIBR) for supporting the research collaboration, providing the 50,000 small molecule drug library and screening facilities. We thank members of the Douglas Auld and Kirk Wright laboratory at NIBR, particularly Zachary Nguyen, for their kind guidance and support with the high throughput screening, technical training, and SPR resources. We also would like to thank all members of the Tenen and Chai groups, particularly Xi Tian and Jing Ping Tang, for their assistance in the experimental studies. Graphical abstract, Fig. 1b,1i, Fig. 6e, Fig. 8, Supplementary Figure 1c, 2d, and 6c were created with BioRender.com.

## References

1. Ou L, Setegne MT, Elliot J, Shen F, and Dassama LM. Protein-based degraders: from chemical biology tools to neo-therapeutics. Chemical Reviews. 2025;125(4):2120–83.

2. Zhang C, Liu Y, Li G, Yang Z, Han C, Sun X, et al. Targeting the undruggables—the power of protein degraders. Science Bulletin. 2024;69(11):1776–97.

3. Moein S, Tenen DG, Amabile G, and Chai L. SALL4: an intriguing therapeutic target in cancer treatment. Cells. 2022;11(16):2601.

4. Yong KJ, Gao C, Lim JS, Yan B, Yang H, Dimitrov T, et al. Oncofetal gene SALL4 in aggressive hepatocellular carcinoma. N Engl J Med. 2013;368(24):2266–76.

5. Liu Y-C, Kwon J, Fabiani E, Xiao Z, Liu YV, Follo MY, et al. Demethylation and Up-Regulation of an Oncogene after Hypomethylating Therapy. New England Journal of Medicine. 2022;386(21):1998–2010.

6. Kwon J, Liu YV, Gao C, Bassal MA, Jones AI, Yang J, et al. Pseudogene-mediated DNA demethylation leads to oncogene activation. Science Advances. 2021;7(40):eabg1695.

7. Zeng SS, Yamashita T, Kondo M, Nio K, Hayashi T, Hara Y, et al. The transcription factor SALL4 regulates stemness of EpCAM-positive hepatocellular carcinoma. J Hepatol. 2014;60(1):127–34.

8. Yong KJ, Li A, Ou WB, Hong CK, Zhao W, Wang F, et al. Targeting SALL4 by entinostat in lung cancer. Oncotarget. 2016;7(46):75425–40.

9. Li A, Jiao Y, Yong KJ, Wang F, Gao C, Yan B, et al. SALL4 is a new target in endometrial cancer. Oncogene. 2015;34(1):63–72.

10. Zhang X, Yuan X, Zhu W, Qian H, and Xu W. SALL4: an emerging cancer biomarker and target. Cancer Lett. 2015;357(1):55–62.

11. Han SX, Wang JL, Guo XJ, He CC, Ying X, Ma JL, et al. Serum SALL4 is a novel prognosis biomarker with tumor recurrence and poor survival of patients in hepatocellular carcinoma. J Immunol Res. 2014;2014:262385.

12. Li A, Yang Y, Gao C, Lu J, Jeong HW, Liu BH, et al. A SALL4/MLL/HOXA9 pathway in murine and human myeloid leukemogenesis. J Clin Invest. 2013;123(10):4195–207.

13. Yuan X, Zhang X, Zhang W, Liang W, Zhang P, Shi H, et al. SALL4 promotes gastric cancer progression through activating CD44 expression. Oncogenesis. 2016;5(11):e268.

14. Matyskiela ME, Couto S, Zheng X, Lu G, Hui J, Stamp K, et al. SALL4 mediates teratogenicity as a thalidomide-dependent cereblon substrate. Nat Chem Biol. 2018;14(10):981–7.

15. Donovan KA, An J, Nowak RP, Yuan JC, Fink EC, Berry BC, et al. Thalidomide promotes degradation of SALL4, a transcription factor implicated in Duane Radial Ray syndrome. Elife. 2018;7.

16. Ma Y, Cui W, Yang J, Qu J, Di C, Amin HM, et al. SALL4, a novel oncogene, is constitutively expressed in human acute myeloid leukemia (AML) and induces AML in transgenic mice. Blood. 2006;108(8):2726–35.

17. Pompili L, Porru M, Caruso C, Biroccio A, and Leonetti C. Patient-derived xenografts: a relevant preclinical model for drug development. J Exp Clin Cancer Res. 2016;35(1):189.

18. Gillespie M, Jassal B, Stephan R, Milacic M, Rothfels K, Senff-Ribeiro A, et al. The reactome pathway knowledgebase 2022. Nucleic acids research. 2022;50(D1):D687–D92.

19. Bonner WM, Redon CE, Dickey JS, Nakamura AJ, Sedelnikova OA, Solier S, et al. γH2AX and cancer. Nature Reviews Cancer. 2008;8(12):957–67.

20. Yang C, Xie S, Wu Y, Ru G, He X, Pan HY, et al. Prognostic implications of cell division cycle protein 45 expression in hepatocellular carcinoma. PeerJ. 2021;9:e10824.

21. Jia W, Xie L, Wang X, Zhang Q, Wei B, Li H, et al. The impact of MCM6 on hepatocellular carcinoma in a Southern Chinese Zhuang population. Biomed Pharmacother. 2020;127:110171.

22. Deng J, Zhong F, Gu W, and Qiu F. Exploration of Prognostic Biomarkers among Replication Factor C Family in the Hepatocellular Carcinoma. Evol Bioinform Online. 2021;17:1176934321994109.

23. Wang M, Xie T, Wu Y, Yin Q, Xie S, Yao Q, et al. Identification of RFC5 as a novel potential prognostic biomarker in lung cancer through bioinformatics analysis. Oncol Lett. 2018;16(4):4201–10.

24. Li W-j, Wu D-w, Zhou Y-f, Zhang C-w, and Liao X-w. Prognostic biomarker replication factor C subunit 5 and its correlation with immune infiltrates in acute myeloid leukemia. Hematology. 2022;27(1):555–64.

25. Zhao X, Wang Y, Li J, Qu F, Fu X, Liu S, et al. RFC2: a prognosis biomarker correlated with the immune signature in diffuse lower-grade gliomas. Scientific Reports. 2022;12(1):3122.

26. Ji Z, Li J, and Wang J. Up-regulated RFC2 predicts unfavorable progression in hepatocellular carcinoma. Hereditas. 2021;158(1):17.

27. Human Protein Atlas.

28. Zheng Ms B, Wang Ms H, Wang Ms JX, Liu Ms ZH, Zhang Md P, and Zhang Md D. The Clinical Significance of RMI2 in Hepatocellular Carcinoma. Technol Cancer Res Treat. 2021;20:15330338211045496.

29. Li Y, He X, Zhang X, Xu Y, Chen W, Liu X, et al. RMI2 is a prognostic biomarker and promotes tumor growth in hepatocellular carcinoma. Clin Exp Med. 2022;22(2):229–43.

30. Palumbo A, Facon T, Sonneveld P, Blade J, Offidani M, Gay F, et al. Thalidomide for treatment of multiple myeloma: 10 years later. *Blood*, The Journal of the American Society of Hematology. 2008;111(8):3968–77.

31. Matyskiela ME, Couto S, Zheng X, Lu G, Hui J, Stamp K, et al. SALL4 mediates teratogenicity as a thalidomide-dependent cereblon substrate. Nature chemical biology. 2018;14(10):981–7.

32. Schuffenhauer A, Schneider N, Hintermann S, Auld D, Blank J, Cotesta S, et al. Evolution of Novartis’ Small Molecule Screening Deck Design. J Med Chem. 2020;63(23):14425–47.

33. Canham SM, Wang Y, Cornett A, Auld DS, Baeschlin DK, Patoor M, et al. Systematic Chemogenetic Library Assembly. Cell Chem Biol. 2020;27(9):1124–9.

34. PubChem Compound Summary for CID 88046120, CID 88046120. National Center for Biotechnology Information. 2023;2023.

35. Lu G, Middleton RE, Sun H, Naniong M, Ott CJ, Mitsiades CS, et al. The myeloma drug lenalidomide promotes the cereblon-dependent destruction of Ikaros proteins. Science. 2014;343(6168):305–9.

36. Martinez Molina D, Jafari R, Ignatushchenko M, Seki T, Larsson EA, Dan C, et al. Monitoring drug target engagement in cells and tissues using the cellular thermal shift assay. Science. 2013;341(6141):84–7.

37. Kalinichenko VV, Major ML, Wang X, Petrovic V, Kuechle J, Yoder HM, et al. Foxm1b transcription factor is essential for development of hepatocellular carcinomas and is negatively regulated by the p19ARF tumor suppressor. Genes Dev. 2004;18(7):830–50.

38. Tamano S, Merlino GT, and Ward JM. Rapid development of hepatic tumors in transforming growth factor alpha transgenic mice associated with increased cell proliferation in precancerous hepatocellular lesions initiated by N-nitrosodiethylamine and promoted by phenobarbital. Carcinogenesis. 1994;15(9):1791–8.

39. Bushnell B. BBTools software packag. e. 2014.

40. Bolger AM, Lohse M, and Usadel B. Trimmomatic: a flexible trimmer for Illumina sequence data. Bioinformatics. 2014;30(15):2114–20.

41. Dobin A, Davis CA, Schlesinger F, Drenkow J, Zaleski C, Jha S, et al. STAR: ultrafast universal RNA-seq aligner. Bioinformatics. 2013;29(1):15–21.

42. Li H, Handsaker B, Wysoker A, Fennell T, Ruan J, Homer N, et al. The Sequence Alignment/Map format and SAMtools. Bioinformatics. 2009;25(16):2078–9.

43. Ramirez F, Dundar F, Diehl S, Gruning BA, and Manke T. deepTools: a flexible platform for exploring deep-sequencing data. Nucleic Acids Res. 2014;42(Web Server issue):W187-91.

44. Liao Y, Smyth GK, and Shi W. featureCounts: an efficient general purpose program for assigning sequence reads to genomic features. Bioinformatics. 2014;30(7):923–30.

45. Kramer A, Green J, Pollard J, Jr., and Tugendreich S. Causal analysis approaches in Ingenuity Pathway Analysis. Bioinformatics. 2014;30(4):523–30.

