## Supplementary figures for "SALLB not targeted by IMiDs is important for maintenance of liver cancer cells"

### Supplementary Data

- **Supplementary Figures**

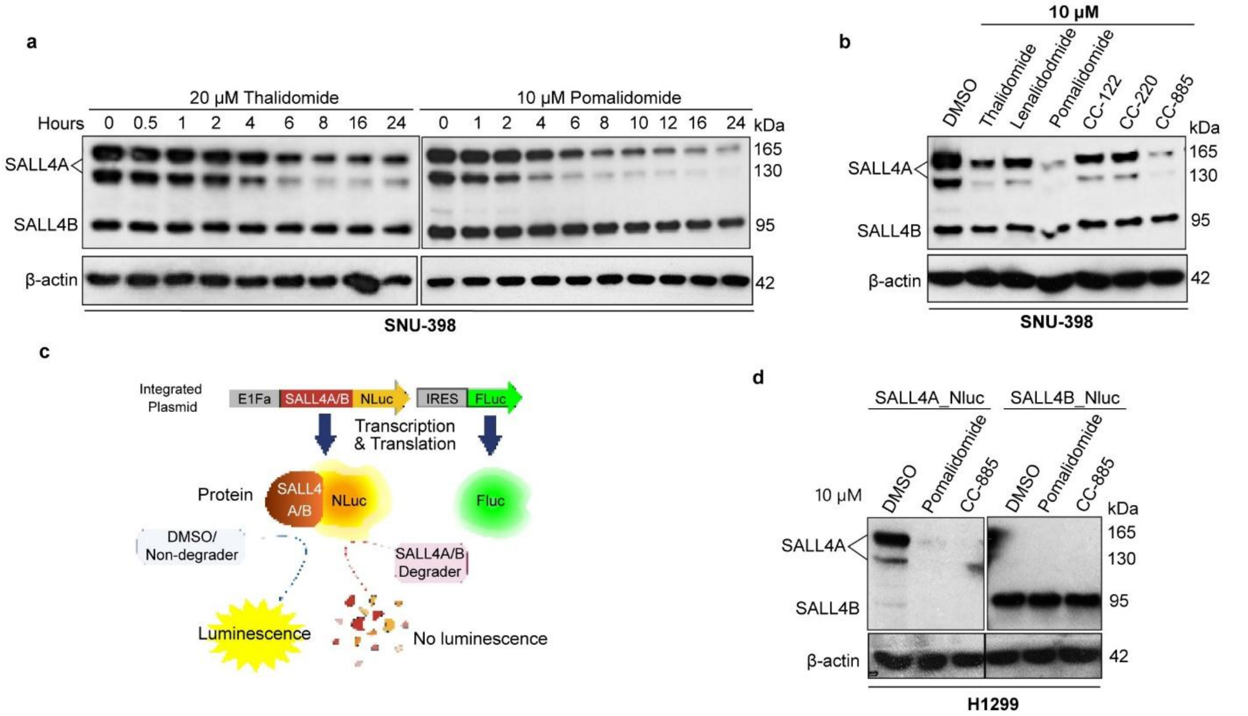


**Supplementary Figure 1 | IMiDs induce degradation of SALL4A, but not SALL4B. a,** Immunoblots showing thalidomide and pomalidomide selectively degraded endogenous SALL4A in SNU-398 cells over time (representative of n = 2). **b,** Immunoblots showing IMiDs degraded endogenous SALL4A, but not SALL4B, in SNU-398 cells after 12 hours treatment (10 μM). **c,** Schematic of dual-luciferase reporter system expressing SALL4A- or SALL4B-Nluc with Fluc as internal control. **d,** Nluc/Fluc ratios after 16 hours IMiD treatment in H1299 cells expressing SALL4A-Nluc or SALL4B-Nluc (n = 2; mean ± SD). i, Immunoblots showing pomalidomide and CC-885 selectively degraded SALL4A-Nluc, but not SALL4B-Nluc, in H1299 cells after 16 h treatment.

.

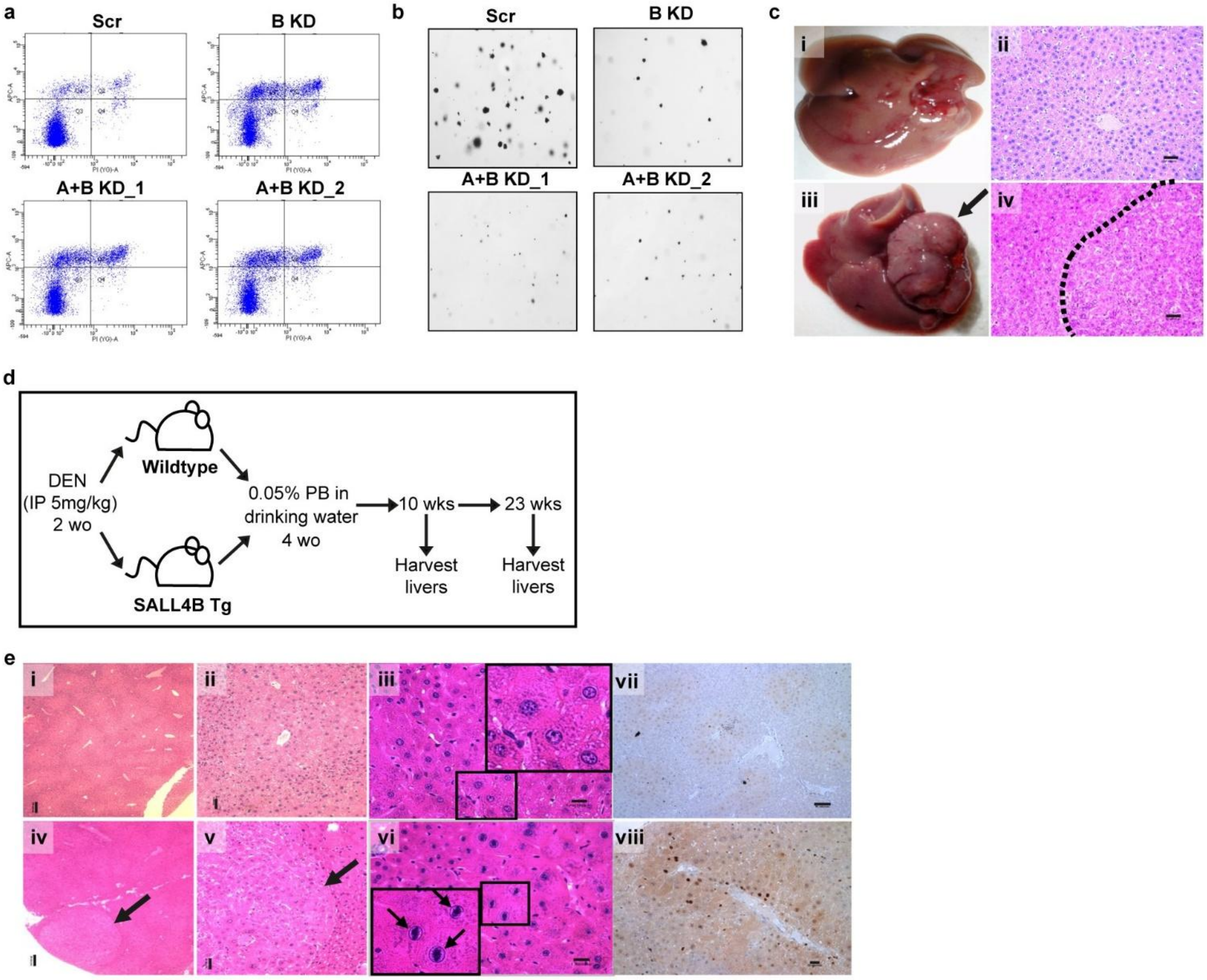


**Supplementary Figure 2 | SALL4-mediated HCC and lung cancer require SALL4B for cell survival. a,** Representative image of Annexin V+ cells analyzed by flow cytometry from SNU398 HCC cells infected with scramble or SALL4 shRNA virus on day 5 post-infection. **b,** Representative image of anchorage-independent soft agar colony formation assay in SNU-398 HCC cells infected with scramble or SALL4 shRNA virus. Bar values represent mean ± SD (n=5). Student's t-test, ****P < 0.0001. **c,** Aged SALL4B transgenic mice developed spontaneous HCC. (i,iii) Gross morphology and (ii,iv) histology following hematoxylin and eosin (H&E) staining of livers of C57BL/6 (i,ii) wild type and (iii,iv) SALL4B transgenic mice. (iii) Arrow indicates liver tumor. (iv) Dashed line indicates normal and tumor (right) boundary. (ii,iv) 200X, bar = 20 µm. **d,** Design of two-stage chemical carcinogenesis experiment. N-nitrosodiethylamine (DEN) and phenobarbital (PB) were used to induce pre-neoplastic hepatic lesions in the wildtype and SALL4B transgenic mice (Tg). Livers were harvested at 10- and 23-experimental weeks (wks). IP: intraperitoneal, wo: weekold. **e,** Young SALL4B transgenic mice were vulnerable to chemical carcinogenesis (DEN/PB). H&E of (i-iii) wildtype and (iv-vi) SALL4B transgenic livers after 23-week DEN/PB exposure. (iv, v) Arrows show cellular alteration foci. (iii) Magnified regions show interphase cells. (vi) Magnified regions and arrows show increased mitosis. Ki-67 immunohistochemistry staining shows increased proliferation in (viii) SALL4B transgenic livers compared to (vii) wild type livers after 23-weeks DEN/PB exposure. (i, iv) 40X, bar = 100 µm. (ii, v) 100X, bar = 40 µm. (iii, vi) 400X, bar = 1 µm.

**a**

| Compound Pubchem ID & SMILES | Thermal Shift Assay | | SALL4B_Luc Reporter Assay |
|---|---|---|---|
| | Type | ΔTm (°C)* | % SALL4B Downregulation† |
| QE (CID: 9796445)<br>COC1=CC=CC=C1NC(=O)N C2=C(C=C(C=C2)[N+](=O)[O-])O | Positive shifter | 1.15 ± 0.01 | 39.8 ± 7.1 |
| RE (CID: 5322399)<br>CC(=O)C1=CC2=C(C=C1)NC(C3C2 C=CC3)C4=CC5=C(C=C4Br)OCO5 | Negative shifter | 0.56 ± 0.08 | 34.6 ± 29.3 |
| CC (CID: 17534)<br>C1=CC=C2C(=C1)C=CC3=C2C=C (C4=CC=CC=C34)N | Positive shifter | 1.52 ± 0.07 | 49.3 ± 8.5 |
| CA (CID: NA)<br>FC(F)(F)c4cccc(CN1CCCn3c(CC1) nc2ccccc23)c4 | Negative shifter | 0.44 ± 0.16 | 54.9 ± 34.1 |
| VB (CID: 4276127)<br>CCN(CC)C(=O)COC(=O)C1=C2CCC (=CC3=CC=C(C=C3)O)C2=NC4=CC=CC=C41 | Positive shifter | 0.20 ± 0.05 | 53.2 ± 1.0 |

* SALL4 (1-300) melting temperature shift, mean ± SD (n = 2). [compound] = 100 μM, 30mins
† Normalized to DMSO control, mean ± SD (n = 3). [compound] = 8 μM, 6h treatment, H1299 cells

**b**

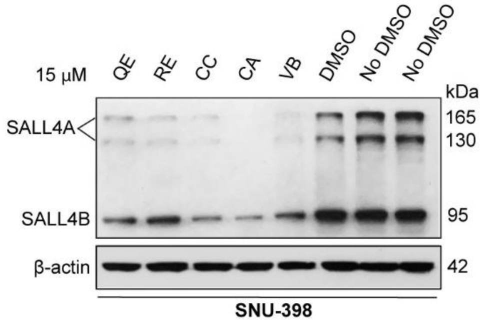


**c**

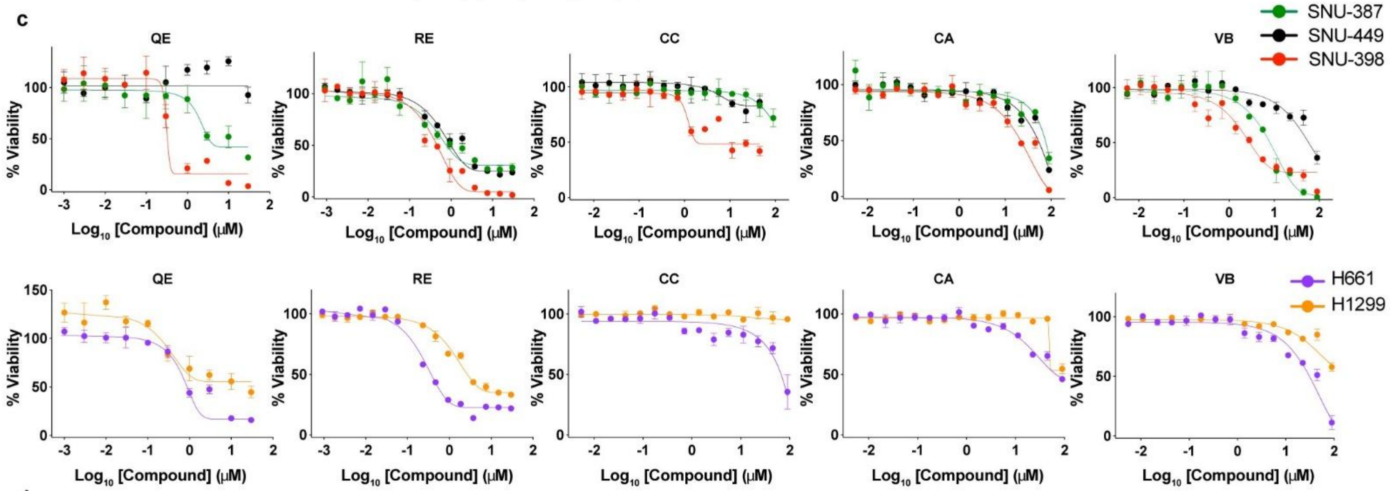


**d**

| Cancer type | Cell line | SALL4B dependency* | 72h $EC_{50}$ (μM) | | | | |
|---|---|---|---|---|---|---|---|
| | | | QE | RE | CC | CA | VB |
| HCC | SNU-398 | dependent | 0.31 | 0.45 | 12.0 | 21.7 | 2.67 |
| HCC | SNU-387 | independent | 3.27 | 0.75 | >90 | 74.8 | 8.02 |
| HCC | SNU-449 | independent | >30 | 0.96 | >90 | 52.5 | 58.2 |
| NSCLC | H661 | dependent | 0.42 | 0.38 | 66.7 | 66.9 | 31.0 |
| NSCLC | H1299 | independent | >30 | 2.67 | >90 | >90 | >90 |

*Cancer cell dependency on SALL4B for survival validated by SALL4B-specific knockdown via shRNA

**Supplementary Figure 3 | Summary of validated screen hits. a,** Table showing summarized data from thermal shift assays and SALL4B dual-luciferase reporter assays of the 5 validated hits. **b,** Immunoblots showing treatment with 15 μM for 16h of the 5 validated hits resulted in downregulation of endogenous SALL4B in SNU-398 HCC cells. **c,** 72h cell viability response curves of the 5 validated screening hits in a panel of SALL4B-dependent versus independent HCC (top) and NSCLC (bottom) cell lines. Data represent triplicates mean ± SD. **d,** Table summarizing 72h $EC_{50}$ values of 5 validated hits in various tumor cell lines from c. HCC: hepatocellular carcinoma, NSCLC: non-small-cell lung carcinoma.

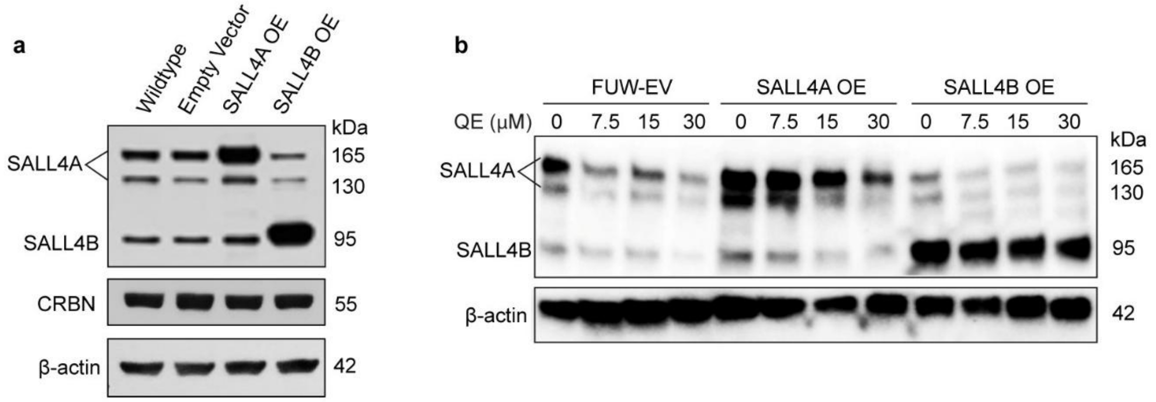


**Supplementary Figure 4 | Rescue of QE therapeutic effects by SALL4 isoform overexpression. a,** Immunoblots showing overexpression of SALL4A or SALL4B in SNU-398 cells transduced with pFUW–SALL4A–mCherry, pFUW–SALL4B–mCherry or empty vector (representative of n = 3). **b,** Immunoblots showing SALL4A or SALL4B levels in SNU-398 cells overexpressing SALL4 isoforms after 24 hours treatment with QE at increasing doses.

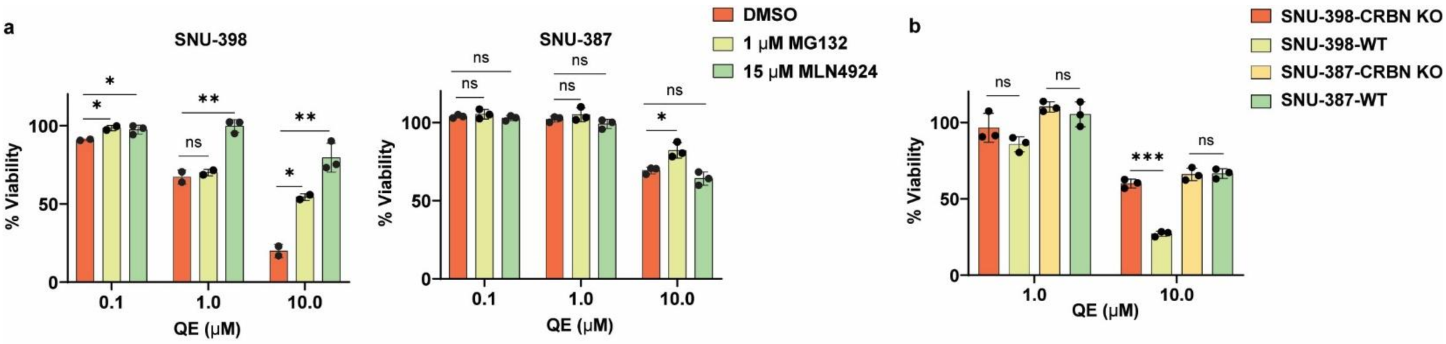


**Supplementary Figure 5| Rescue of QE therapeutic effects by blockade of CRBN activity. a,** Viability of SNU-398 and SNU-387 cells after 72 hours co-treatment with QE (10 μM) and MG132 (1 μM) or MLN4924 (15 μM) (n = 3; mean ± SD). **b** Viability of SNU-398 wild-type, SNU-398 CRBN–KO, SNU-387 wild-type and SNU-387 CRBN–KO cells after 72 hours QE treatment (n = 3; mean ± SD).

**Statistics:** One-way ANOVA with Dunnett's post hoc test versus DMSO at each dose for (c). Unpaired two-tailed t-test for (d). $^{*}P < 0.05$, $^{**}P < 0.01$, $^{***}P < 0.001$, $^{****}P < 0.0001$; ns, not significant ($P \geq 0.05$).

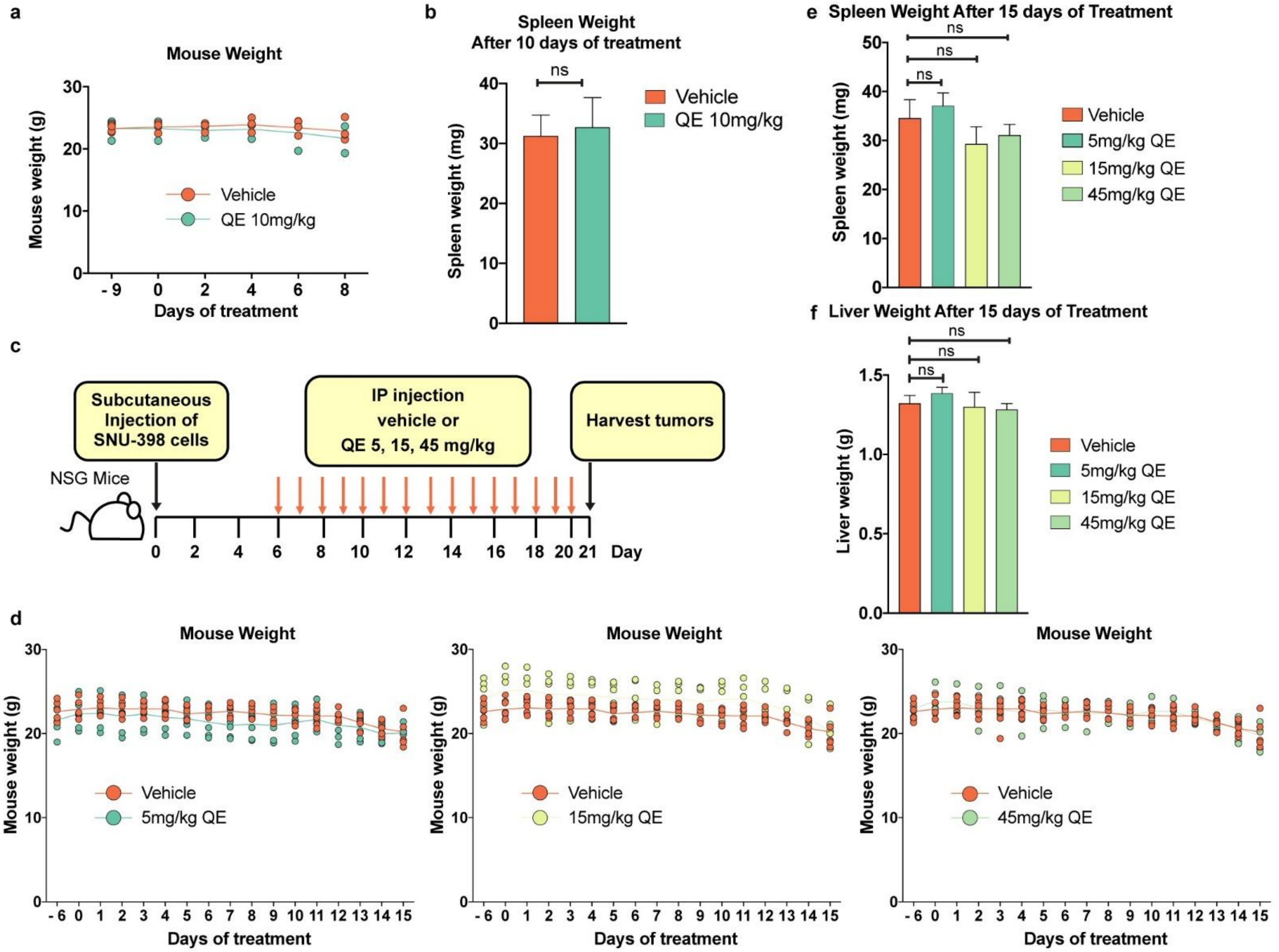


**Supplementary Figure 6 | In vivo anti-tumour tolerability and dose expansion of QE. a,** Body weight of mice treated with vehicle (n = 4) or QE (10 mg/kg, q.d.; n = 3) for 10 days; each data point represents an individual mouse. **b,** Spleen weights at day 10 of the pilot study (n = 3-4; mean ± SEM). **c,** Schematic of dose-expansion study using SNU-398 xenografts. **d,** Body weight of mice treated with vehicle or QE at 5, 15 or 45 mg/kg (q.d.) for 15 days (n = 6 per group); each data point represents an individual mouse. **e,** Spleen weights at day 15 (n = 6; mean ± SEM). **f,** Liver weights at day 15 (n = 6; mean ± SEM). **Statistics:** Unpaired two-tailed t-test for (b). One-way ANOVA with Dunnett's post hoc test versus vehicle for (e) and (f). ns, not significant ($P \geq 0.05$).

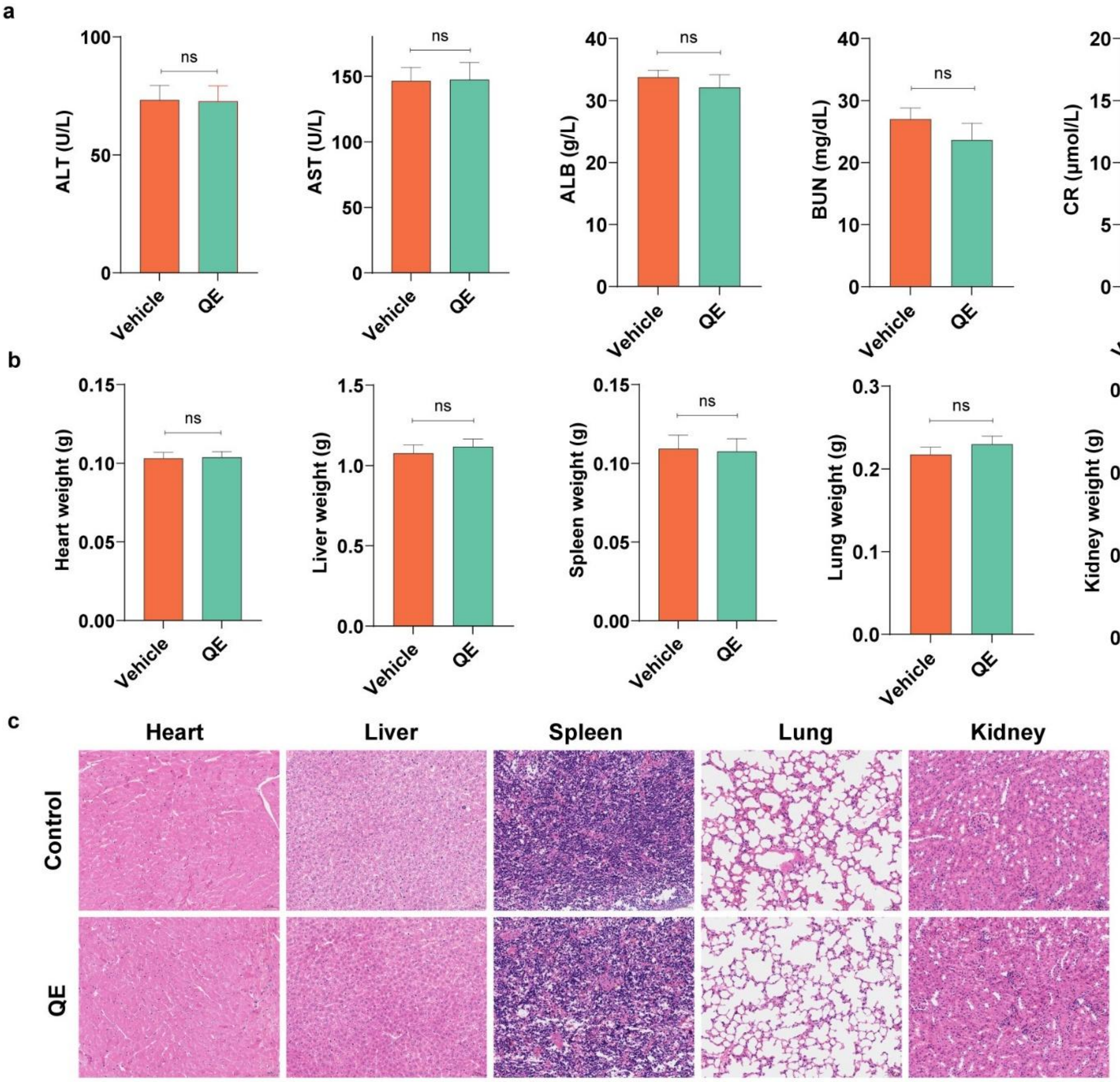


**d** Blood Routine Analysis

| | Control | QE |
|---|---|---|
| WBC（10^9/L） | 8.5±1.97 | 5.8±1.15 |
| RBC（10^12/L） | 8.3±1.1 | 9.1±1.5 |
| HGB（g/L） | 120.0±13.7 | 118.5±13.3 |
| HCT（%） | 40.6±4.5 | 39.3±4.2 |
| MCV（fL） | 52.0±3.1 | 52.3±5.5 |
| PCT（%） | 0.509±0.087 | 0.546±0.109 |

WBC = White Blood Cell; RBC = Red Blood Cell; HGB= Haemoglobin; HCT= Haematocrit; MCV =Mean Corpuscular Volume; PCT = Procalcitonin

**Supplementary Figure 7. Antitumor activity of QE in SALL4-high HCC PDX model.** **a,** Serum biochemistry (ALT, AST, ALB, BUN and creatinine) on day 39 in vehicle- and QE-treated mice (n = 8; mean ± SEM). **b,** Organ weights (heart, liver, spleen, lung and kidney) from vehicle- and QE-treated mice on day 39 (n = 14; mean ± SEM). **c,** H&E staining of major organs from vehicle- and QE-treated mice (scale bar, 50 μm). **d,** Peripheral blood counts (WBC, RBC, HGB, HCT, MCV and PCT) on day 39 in vehicle- and QE-treated mice (n = 8; mean ± SEM).
**Statistics:** Unpaired two-tailed t-test for (b) and (c). ns, not significant ($P \geq 0.05$).

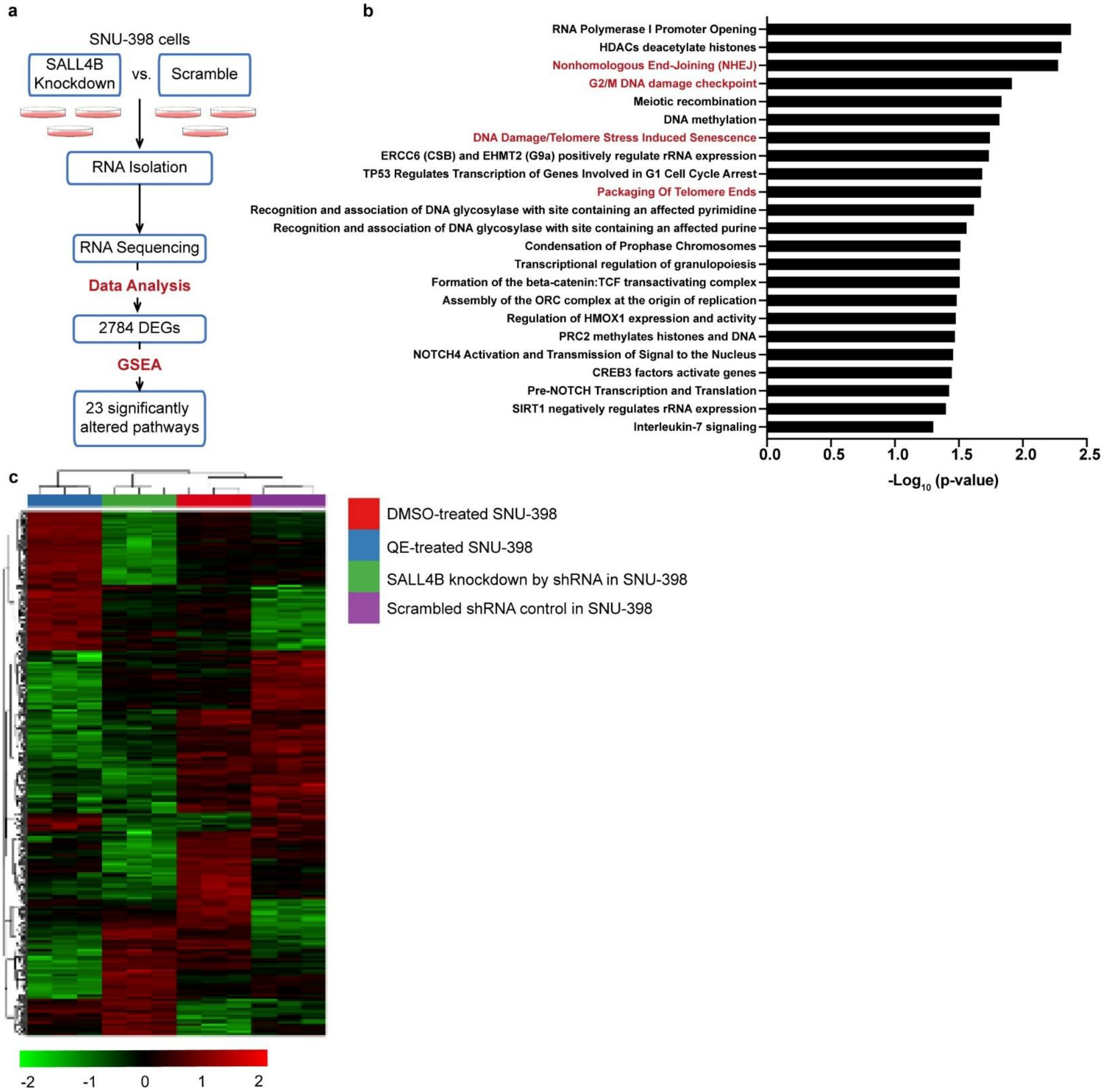


**Supplementary Figure 8 | SALL4B knockdown and QE treatment induce convergent transcriptional repression of DNA repair and replication pathways in SALL4-dependent HCC cells. a,** RNA-seq analysis of SNU-398 cells 48 h after transduction with scramble or SALL4B shRNA. Differentially expressed genes (DEGs; $\log_2 FC > 0.5$, FDR < 0.05) were subjected to GSEA. **b,** Significantly enriched pathways altered by SALL4B knockdown in SNU-398 cells (n = 3; shown as $-\log_{10}$ P values). **c,** Hierarchical clustering heat map of gene expression changes in SNU-398 cells following SALL4B knockdown or QE treatment (10 μM, 24 hours), relative to scramble or DMSO controls. Each column represents an individual sample and each row a gene.
**Statistics:** Unpaired two-tailed t-test for (d). ns, not significant (P ≥ 0.05).
